# Optimal hearing aid design through restoration of the neural code

**DOI:** 10.64898/2026.02.02.703273

**Authors:** Fotios Drakopoulos, Lloyd Pellatt, Shievanie Sabesan, Yiqing Xia, Tingchen Gong, Andreas Fragner, Nicholas A Lesica

## Abstract

Hearing loss introduces complex distortions in the neural coding of sound that current hearing aids fail to address. Here, we combine electrophysiology and deep learning to identify novel sound processing strategies to correct these distortions. We use large-scale intracranial recordings from the gerbil inferior colliculus to train deep neural network models of neural coding (ICNets) to serve as in silico surrogates for brains with normal and impaired hearing. We then use the ICNets to train another network (AidNet) to act as an optimal hearing aid, providing the individualized sound processing required to elicit normal neural activity in impaired brains. We find that AidNet outperforms state-of-the-art hearing aid processing by a wide margin in correcting distortions in neural coding and deficits in simulated phoneme recognition. Much of this advantage was retained when AidNet’s parameters were swapped across animals with similar hearing thresholds, suggesting that AidNet can compensate for complex effects of hearing loss without direct access to neural recordings. These results demonstrate that closed-loop optimization can identify novel and generalizable strategies for neural restoration, providing a foundation for the development of more effective hearing aids as well as other sensory devices and neurotechnologies.

## Introduction

Hearing loss affects more than a billion people globally, causing widespread disability [1] and declines in mental health and wellbeing [2]. For the most common forms of hearing loss, the only treatment available is a hearing aid. Traditional hearing aids are designed to provide amplification to restore lost sensitivity as measured by an audiogram, which specifies hearing thresholds at different frequencies. This approach has two key shortcomings: The use of constrained signal processing and the focus on thresholds, both of which limit the potential to compensate for the complex effects of hearing loss on auditory processing [3]. While most devices employ additional features (including AI-based) to address specific listening scenarios [4, 5], their real-world utility for sounds such as speech in complex background noise [6, 7] or music [8, 9] remains limited.

We have taken a new approach to hearing aid design, operating from the perspective of the neural code — the brain activity patterns that underlie perception. If hearing is impaired, it is because the details of the neural code have become distorted. Thus, to restore perception to normal, a hearing aid should process sound as needed to elicit neural activity in an impaired auditory system that matches the activity elicited by the original sound in a normal auditory system [10–12]. From this perspective, the key challenge in hearing aid design is identifying the sound processing required to correct the distortions in the brain’s internal representation of the full auditory scene.

Compensating for the complex effects of hearing loss on neural coding, which are only partly understood, is beyond the scope of classical engineering or analytical approaches. Fortunately, deep learning provides tools for neural control [13–15] that enable data-driven identification of the sensory inputs required to elicit desired activity patterns. By allowing for flexible signal processing and considering the effects of hearing loss on the full neural code, these tools have the potential to address the key shortcomings of current devices. We demonstrate how these tools can be used to optimize hearing aid processing in a closed-loop framework, iteratively updating the sound processing performed by a neural network until the sound it produces elicits normal activity in an impaired brain.

## Results

To identify hearing aid processing that can compensate for both cochlear damage and downstream plasticity, we focused on the neural code in the inferior colliculus (IC), the midbrain hub of the central auditory pathway. The IC is an obligatory bottleneck where multiple brainstem inputs converge to create a full neural representation of the auditory scene. It is late enough in the auditory pathway to capture many of the plastic changes that follow hearing loss [16], but also early enough that its neural activity is still driven primarily by raw acoustics.

The features of the neural code in the IC that carry information about complex sounds such as speech and music are only observable through high-resolution invasive methods [17, 18], which necessitates the use of an animal model. We used gerbils, a common model for studies of human hearing [19], and made intracranial recordings using silicon probe electrode arrays with hundreds of recording channels [20]. These electrode arrays allow us to observe the timing of action potentials from large populations of neurons, giving us a comprehensive view of the neural code in individual brains and providing the rich datasets required for deep learning.

Our approach to developing optimal hearing aids (AidNets) involves multiple steps (Fig. 1a). We first use IC recordings to train forward models of neural coding (ICNets) that take a sound waveform as input and produce as output the spike patterns that we observe with and without hearing loss. These models are highly accurate and can serve as in silico surrogates for the normal and impaired auditory systems that they are trained to mimic. We then use the normal and impaired ICNets to train AidNets that process sounds as required to restore distorted neural coding to normal, and compare the performance of the trained AidNets against that of traditional hearing aids.

**Fig. 1.**
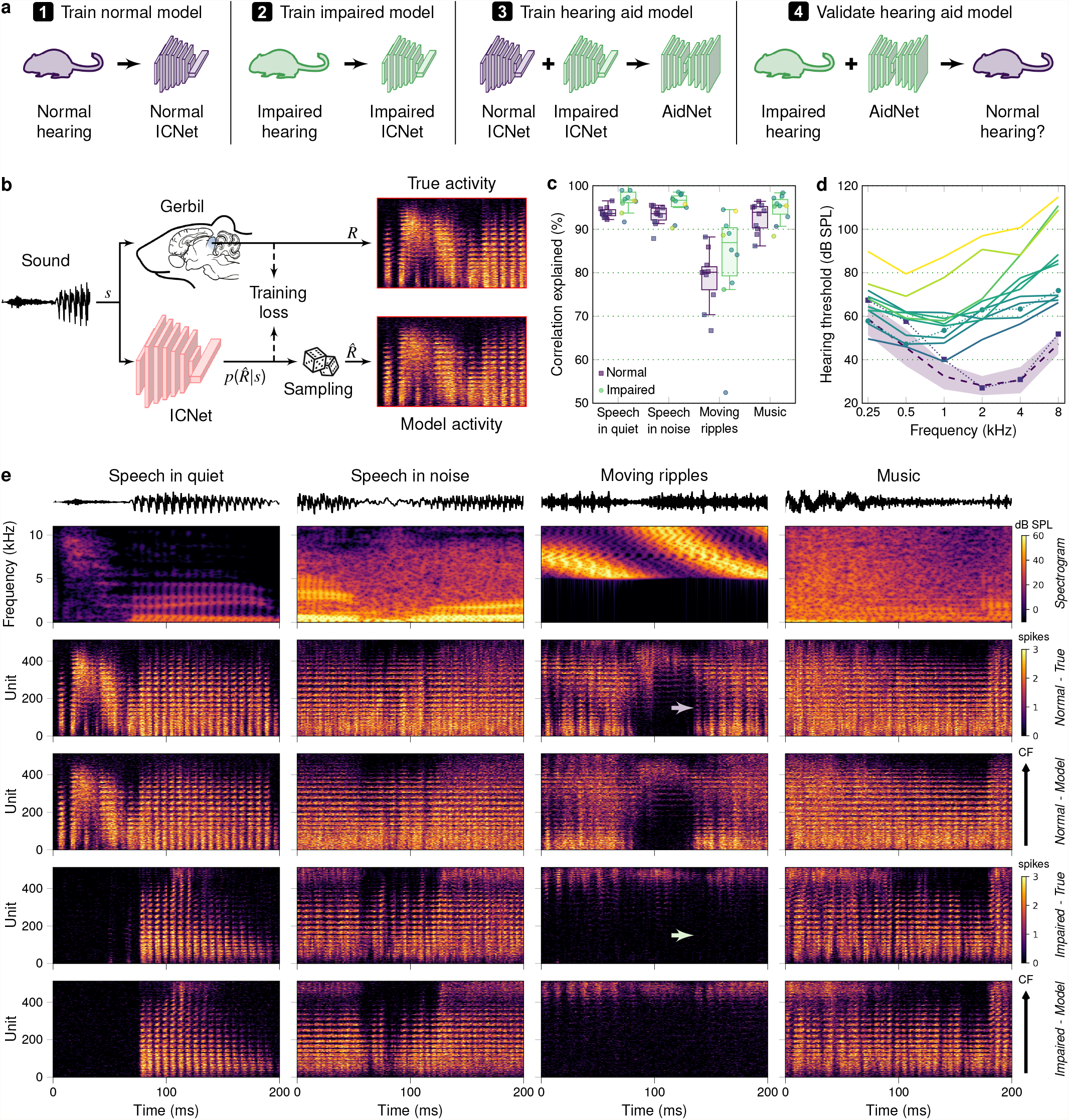
ICNets provide accurate simulation of normal and impaired neural coding. **a**. Schematic diagram of our framework for optimal hearing aid design. **b**. Schematic diagram of our approach to training an ICNet that maps a sound waveform to the neural activity recorded from a single animal. **c**. Predictive power of ICNet models for 10 normal hearing (NH) and 10 hearing impaired (HI) brains. Neural responses to four sounds were simulated by sampling from the distribution 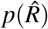 produced by each model. Performance was measured as the percentage of explainable correlation explained by comparing the correlation between simulated and real responses to that between real responses on successive trials. Correlation was computed after flattening the responses by collapsing across time and units. Each dot represents the performance for one animal, and the box plots illustrate the distribution across animals. The horizontal line indicates the median, the box spans the 25th to 75th percentiles (IQR), and the whiskers extend to the most extreme data points within 1.5 times the IQR from the box edges, if any exist. **d**. Hearing thresholds of the HI animals. Thresholds were computed from neural responses to pure tones of different frequencies. The colors indicate the average thresholds for frequencies between 1 and 8 kHz. The dashed line and shaded region indicate the mean and standard deviation of the hearing thresholds of 12 NH animals. The thresholds corresponding to the example responses shown in **e** (and in Figs. 2-3) are indicated by the dotted lines and symbols. **e**. Real and simulated MUA for different sounds. The top two rows show example sound waveforms and spectrograms. The third and fourth rows show recorded and simulated activity for an example NH brain. The fifth and six rows show recorded and simulated activity for an example HI brain. The neural units are organized according to their characteristic frequency (CF; range 0.3-16 kHz). The purple arrow highlights an example of the non-linearity of neural coding: While the sound contains frequencies only above 5 kHz, it elicits strong activity across the full extent of the IC, with low-CF units driven by distortions created by outer hair cell interactions in the healthy cochlea. The green arrow highlights the challenge involved in hearing aid design: After hearing loss, activity in low-CF units is eliminated and cannot be restored by simple amplification because the outer hair cells no longer create distortions. Instead, a hearing aid must include its own non-linear processing that introduces low frequencies into the sound to drive these units directly.

### ICNets provide accurate simulation of neural coding with and without hearing loss

The ultimate success of the AidNet optimization depends on the accuracy of the ICNet models across the full space of sounds that is explored during the optimization process. To provide suitable training datasets, we recorded activity under anesthesia in response to a wide range of sounds, including more than 10 hours of natural and processed speech, music, environmental noises and artificial sounds. We then extracted the multi-unit activity (MUA) from each of 512 electrode channels (from one 256-channel array in the central nucleus of the IC in each hemisphere) and represented it as spike counts in 1.3 ms time bins.

The ICNets are trained to simulate neural activity that is as close as possible to the real neural activity recorded from individual brains [21]. We frame the simulation of neural coding as a classification problem: Given the recent history of sound input, the models are trained using a crossentropy loss to produce an estimate of the full probability distribution of the activity for each neural unit in each time bin, with classes corresponding to the possible spike count values (0-4). The ICNets use a convolutional encoder-decoder architecture, with the encoder mapping the sound waveform to latent features and the decoder mapping the latent features to spike count probabilities for each neural unit and time bin (Fig. 1b).

We first trained ICNets for a collection of normal hearing (NH) brains to serve as potential targets for hearing aid optimization. As an overall measure of ICNet accuracy, we computed the fraction of the explainable correlation (i.e., the correlation of recorded responses on successive trials) that ICNet explained for sounds that were not included in training. The ICNets performed well across all test sounds (Fig. 1c; each purple square represents performance across all units for one NH brain) explaining 93.6% (91.9-96.7) of the explainable correlation for speech in quiet (median and 95% confidence intervals over bootstrap samples), 93.6% (87.3-95.8) for speech in noise, 93.9% (85.6-96.6) for music, and 80.0% (66.1-88.5) for moving ripples.

To assess whether ICNets can also replicate neural coding in hearing impaired (HI) brains, we used noise exposure [22] to introduce varying degrees of hearing loss to a number of animals (Fig. 1d). After the hearing loss stabilized (4-8 weeks after noise exposure), we recorded the training dataset from each animal and trained an ICNet for each individual HI brain. These ICNets also performed well – better, often, than for normal hearing (Fig. 1c; yellow and green circles), explaining 96.7% (91.3-100.5) of the correlation for speech in quiet, 96.6% (89.9-98.7) for speech in noise, 95.4% (88.1-98.5) for music, and 87.2% (51.8-94.8) for moving ripples. We found no correlation between degree of hearing loss and ICNet performance across the four sounds (Spearman’s *r* = -0.0068, *p* = 0.96; hearing loss defined as average threshold as in Fig. 1d).

### Aligning neural activity without correcting the effects of hearing loss

The optimal hearing aid for a given impaired brain would transform sound so that the difference between the impaired brain’s response to the transformed sound and a normal brain’s response to the original sound is minimized. But even without hearing loss, activity recorded on the same electrode channel in different brains will differ (e.g., because of variations in anatomy or electrode placement), creating an alignment problem that complicates cross-brain comparisons (Fig. 2a).

**Fig. 2.**
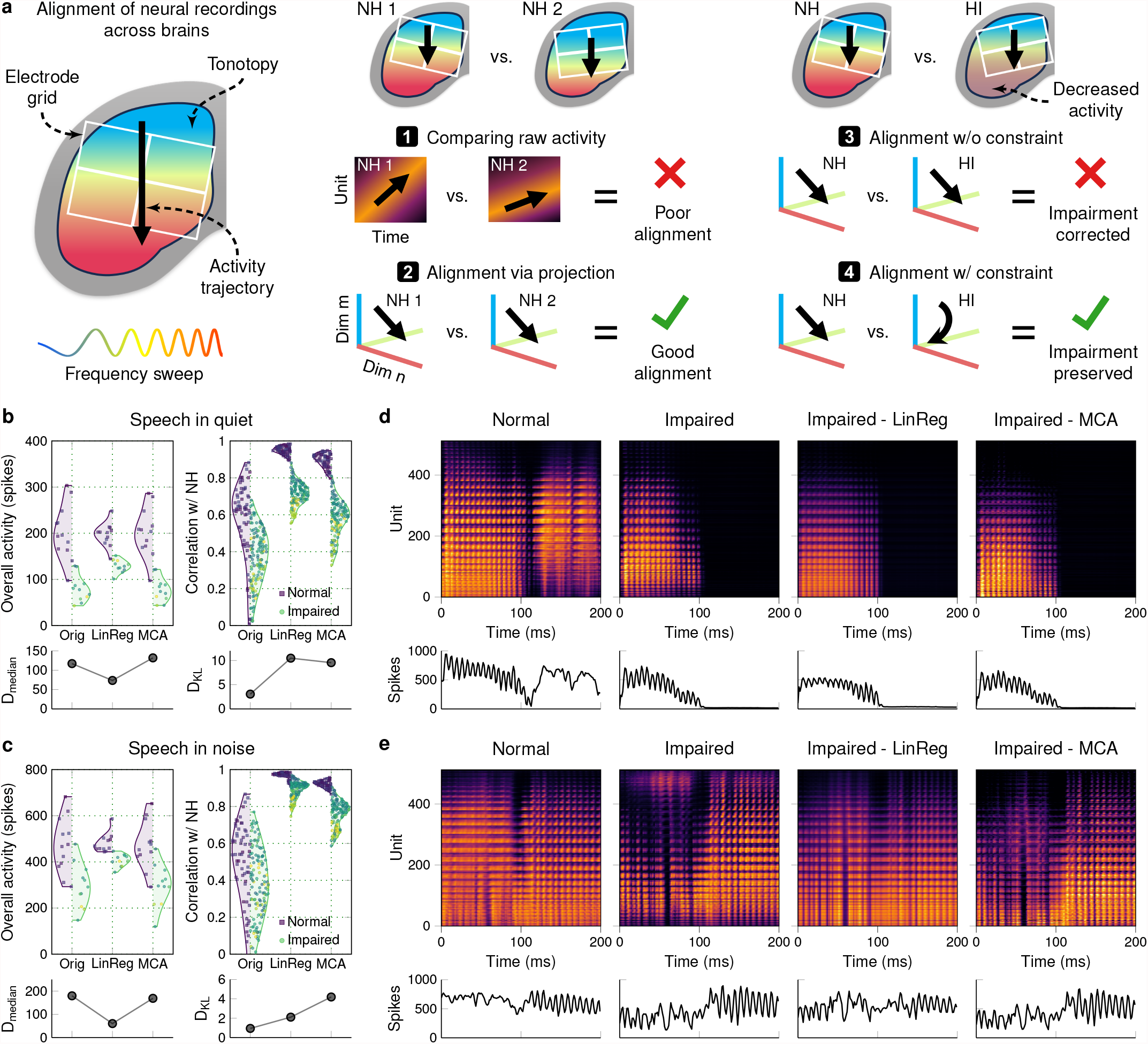
Constrained alignment of recordings across brains preserves the effects of hearing loss. **a**. Schematic diagram illustrating the challenge of aligning neural recordings from brains with and without hearing loss. A tone with increasing frequency elicits activity that travels across the tonotopic gradient in the IC and, consequently, across the electrode grid. (1) Because anatomy and electrode placement vary across brains, raw recordings are idiosyncratic and poorly aligned; (2) Recording idiosyncrasies can be removed via projection into a common space; (3) With hearing loss, activity at the high-frequency end of the tonotopic gradient is decreased, but unconstrained alignment removes this effect; (4) Constrained alignment can remove recording idiosyncasies while preserving effects of hearing loss. **b**. Responses to speech in quiet presented at 50 dB SPL were used to compare the similarity of activity from all NH and HI brains, both before and after alignment. To eliminate neural noise, ICNet simulations were used and the expectation over counts of 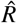 was taken. The top left panel shows total activity (summed across units, averaged over time bins) for each of 12 NH (squares) and 11 HI (circles) brains before and after alignment. The top right panel shows the correlation of activity patterns (after flattening across units) between pairs of NH brains (squares) and between pairs of NH and HI brains (circles). The color code for hearing thresholds is the same as in Fig. 1d. The bottom left panel shows the difference in the medians of the NH and HI activity distributions. The bottom right panel shows the KL divergence between the NH and NH-HI correlation distributions. **c**. Results for responses to speech presented at 65 dB SPL and mixed with restaurant noise at 0 dB SNR, presented as in **b. d**. Example responses to speech in quiet (the syllable “esh”). The HI responses are shown before and after alignment. The top row shows heatmaps with neural activity clipped between 0 and 4. The bottom row shows the activity summed across units. **e**. Example responses to speech in noise (the syllable “shu”), presented as in **d**.

These differences across brains can manifest as variations in both overall activity level and in the correlation of activity patterns between units on matched channels. As shown in the distributions labeled ‘Orig’ in Fig. 2b-c, even across NH brains, the overall activity in response to the same sound can vary by up to a factor of 3 (each purple square represents the summed activity across all units in one NH brain) and the correlation between units on matched channels can range between 0 and 0.9 (each purple square represents the correlation across all matched units for one pair of NH brains). Because of this variation, the distributions of correlations for NH-HI pairs (yellow and green circles) are largely overlapping with the distributions for NH pairs, masking the distortions in neural activity caused by hearing loss.

To ensure that hearing aid optimization is focused on the relevant features, the neural activity from normal and impaired brains must be aligned so that only the differences related to hearing loss remain. Alignment of neural recordings is traditionally done with techniques such as linear regression or canonical-correlation analysis (CCA) to identify transformations that minimize differences in activity [23, 24]. Such techniques may be effective for normal brains, but cannot be used to align normal and impaired brains for our purposes, as they may remove effects of hearing loss. This was evident when we used linear regression to align our IC recordings, as it increased the overall activity in HI brains after alignment (‘LinReg’ activity distributions in Fig. 2b-c).

As alternatives, we tested other alignment methods with different magnitude-preserving constraints: Orthogonal-Rotational Procrustes, Optimal Transport, and Maximum Covariance Analysis (MCA). When tested on activity from an analytical model in which misalignment and hearing loss could be independently controlled, we found that all three methods were effective in correcting misalignment without correcting hearing loss (Supplementary Fig. 1). For our IC recordings, it is impossible to determine directly the degree to which an alignment method is correcting hearing loss. We reasoned that the best alignment method for our purposes would be that which maximizes the separation between the distributions of correlations for NH pairs and NH-HI pairs (i.e., that which minimizes the differences between NH pairs by correcting misalignment while maximizing the differences between NH-HI pairs by preserving the effects of hearing loss). To assess the different methods on this basis, we measured the KL divergence between the correlation distributions for NH pairs and NH-HI pairs after alignment. For speech presented in quiet and in noise, the correlation distributions that were highly overlapping before alignment were well separated after alignment by all three constrained methods (Fig. 2b-c for MCA, Supplementary Fig. 2 for others). Because MCA performed slightly better than the other methods according to the KL divergence metric (and also had other desirable properties, see Supplementary Fig. 1), we chose to move forward with this method for hearing aid optimization.

### AidNets correct distorted neural coding after hearing loss

To train AidNets to act as optimal hearing aids, we used the closed-loop optimization framework illustrated in Fig. 3a. In the normal branch (top), sound was passed through a NH ICNet (one of 9 chosen randomly on each forward pass) to generate a target response. In the impaired branch (bottom), the same sound was passed first through the AidNet and then through a fixed HI ICNet to generate a response to the AidNet-processed sound. AidNet was trained to minimize the differences between the responses from each branch (after alignment with MCA) using a composite loss function that included several neural metrics (see Supplementary Fig. 3) as well as other terms focused on the sound processing itself [11]. Inasmuch as even a truly optimal AidNet will not be able to correct all of the distortions in the neural code, it is important that the training encourages the correction of those distortions that matter most. To focus the optimization on features of neural activity that are likely to carry perceptually relevant information, we also included in the training framework an automatic speech recognition (ASR) model that decoded phonemes from neural activity. The responses from the normal and impaired branches were passed through the phoneme decoder and the difference between the inferred phoneme probability distributions (measured as crossentropy) was included in the AidNet training loss (Supplementary Fig. 4).

**Fig. 3.**
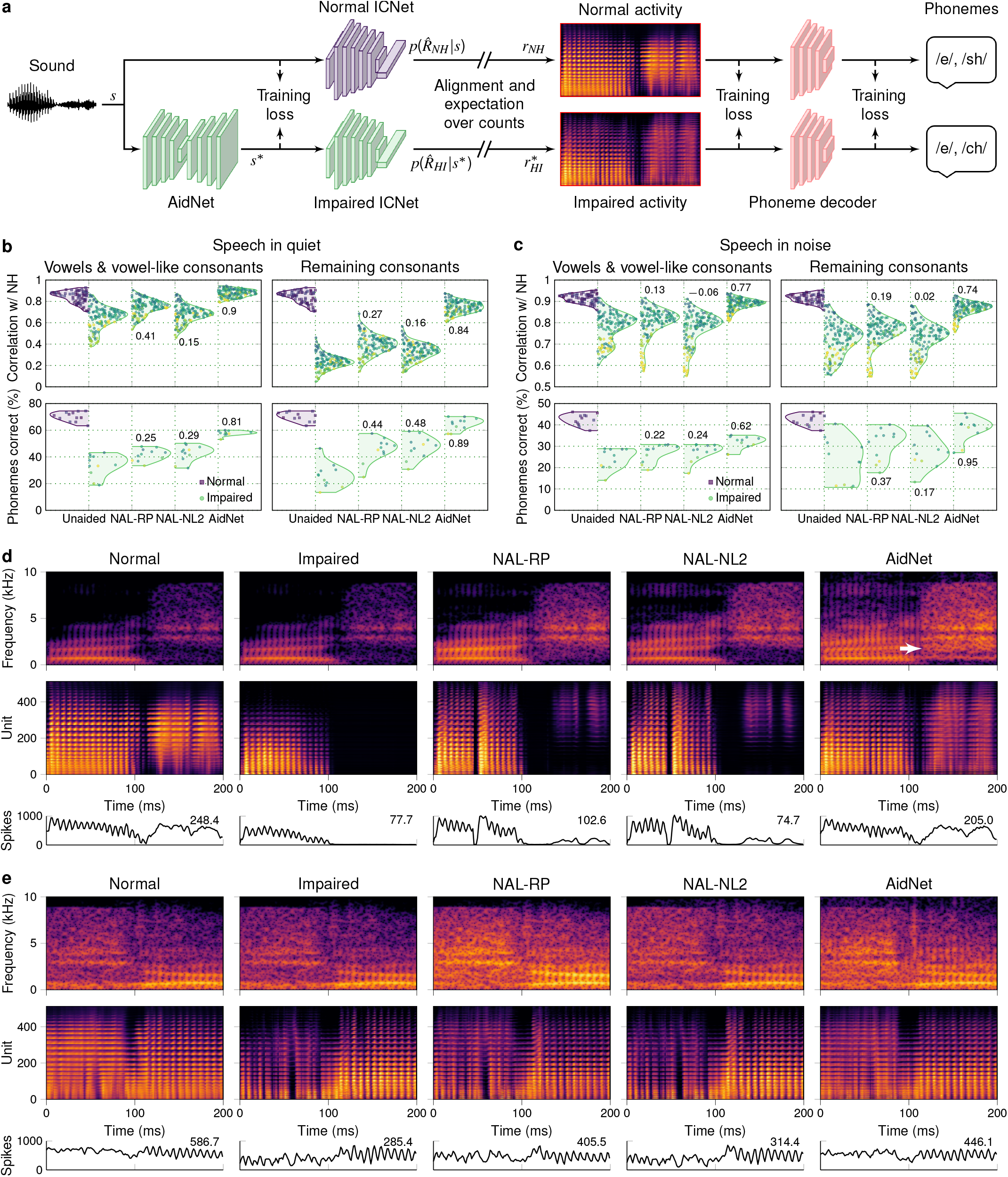
AidNets outperform the state of the art in silico. **a**. Schematic diagram of the AidNet training framework. Only the AidNet was updated during training; the ICNet and phoneme decoder models were fixed. **b**. Restoration results for simulated responses to speech in quiet at 50 dB SPL. For each HI animal, the speech was presented without any processing as well as after processing with the two benchmark amplification strategies (with fitting based on hearing thresholds) and an individually optimized AidNet. To eliminate neural noise, the expectation over counts of 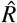 was taken for each ICNet model. The top panels show the correlation between activity patterns from pairs of 12 NH brains (squares), as well as from pairs of 12 NH and 11 HI brains (circles) for each of the indicated processing conditions. Correlation was computed after flattening the responses. The results are shown separately for vowels and vowel-like consonants and the remaining consonants. The bottom panels show the results of phoneme recognition simulated from the responses using the phoneme decoder. Note that the phoneme decoder was trained only on responses to unprocessed sounds from a subset of animals (see Methods and Supplementary Fig. 4). The color code for hearing thresholds is the same as in Fig. 1d. The numbers indicate the average restoration index (for full statistics, see Supplementary Table 1). **c**. Restoration results for speech in noise at 65 dB SPL and 0 dB SNR, presented as in **b. d**. Example spectrograms and responses to speech in quiet (the syllable “esh”). The HI responses are shown both before and after the indicated processing. The bottom row shows the activity summed across units, with the numbers indicating average over time. The white arrow highlights an example of the difference between AidNet and the benchmark amplification strategies: The benchmark strategies increase the intensity of high frequencies, which restores some of the missing activity in high-CF units. AidNet, in addition to amplifying high frequencies, also injects energy at low frequencies, creating additional activity in both low- and high-CF units, with the latter receiving additional drive through the spread of excitation on the cochlea. **e**. Example spectrograms and responses to speech in noise (the syllable “shu”), presented as in **d**.

To evaluate performance after training, we measured the similarity of responses to speech from NH and HI ICNets with and without AidNet processing. We used correlation as our main neural metric and computed results separately across two different phoneme classes: (1) vowels and vowel-like consonants with predominantly low frequencies and harmonic structure and (2) other consonants with higher frequencies for which the loss of sensitivity due to hearing loss is expected to have a more detrimental effect. As benchmarks, we compared the performance of AidNet against linear amplification (NAL-RP) as well as against state-of-the-art multi-channel wide dynamic range compression (WDRC) processing (NAL-NL2), both fit using the hearing thresholds measured for each HI animal (Fig. 1d).

To summarize the restoration achieved by each form of hearing aid processing, we computed a restoration index (*ρ*_*NH,HA*_ −*ρ*_*NH,HI*_)*/*(*ρ*_*NH*_ −*ρ*_*NH,HI*_). The numerator is the difference in the correlations for NH-HI pairs with and without hearing aid (HA) processing, and the denominator is the difference in the correlations for NH and NH-HI pairs. A value of 1 indicates full restoration and a value of 0 indicates that the hearing aid processing provided no benefit. As shown in the top panels of Fig. 3b-c, the benchmark amplification strategies provided only limited benefit (mainly for speech in quiet), with linear amplification performing slightly better than WDRC (overall restoration of 0.25 and 0.07 averaged across all pairs and both phoneme classes for NAL-RP and NAL-NL2, respectively; see Supplementary Table 1 for full statistics). This is consistent with previous results showing that WDRC processing fails to correct distortions in the central neural coding of speech [20]. In contrast, AidNet corrected most of the distortions in neural activity for both speech in quiet and speech in noise (average overall restoration of 0.81) without necessarily providing more amplification (Supplementary Fig. 5). The benefit of AidNet relative to the benchmark processing strategies was also evident when performance was assessed with several other neural metrics (see Supplementary Fig. 3).

We also assessed the degree to which hearing aid processing restored simulated phoneme recognition to normal. We used the same phoneme decoder that was included in AidNet training to decode phonemes from the NH and HI ICNet responses to speech, with and without the different forms of processing. We computed a restoration index based on the percentage of phonemes correctly recognized (*PC*_*HA*_ −*PC*_*HI*_)*/*(*PC*_*NH*_ −*PC*_*HI*_) and found that AidNet was highly effective in restoring simulated phoneme recognition, while the other processing strategies again provided only limited benefit (average overall restoration of 0.34, 0.32 and 0.77 for NAL-RP, NAL-NL2 and AidNet, respectively; see Supplementary Table 1 for full statistics).

### AidNets correct distorted neural coding in vivo

To assess whether the benefits of AidNet were evident in real brains, we designed an experimental protocol that allowed us to train and validate an AidNet within a single recording session (Fig. 4a). For each of three impaired animals, we recorded an abridged training dataset; trained an ICNet; trained an AidNet; and then recorded responses to AidNet-processed sounds as well as sounds processed by the benchmark amplification strategies.

**Fig. 4.**
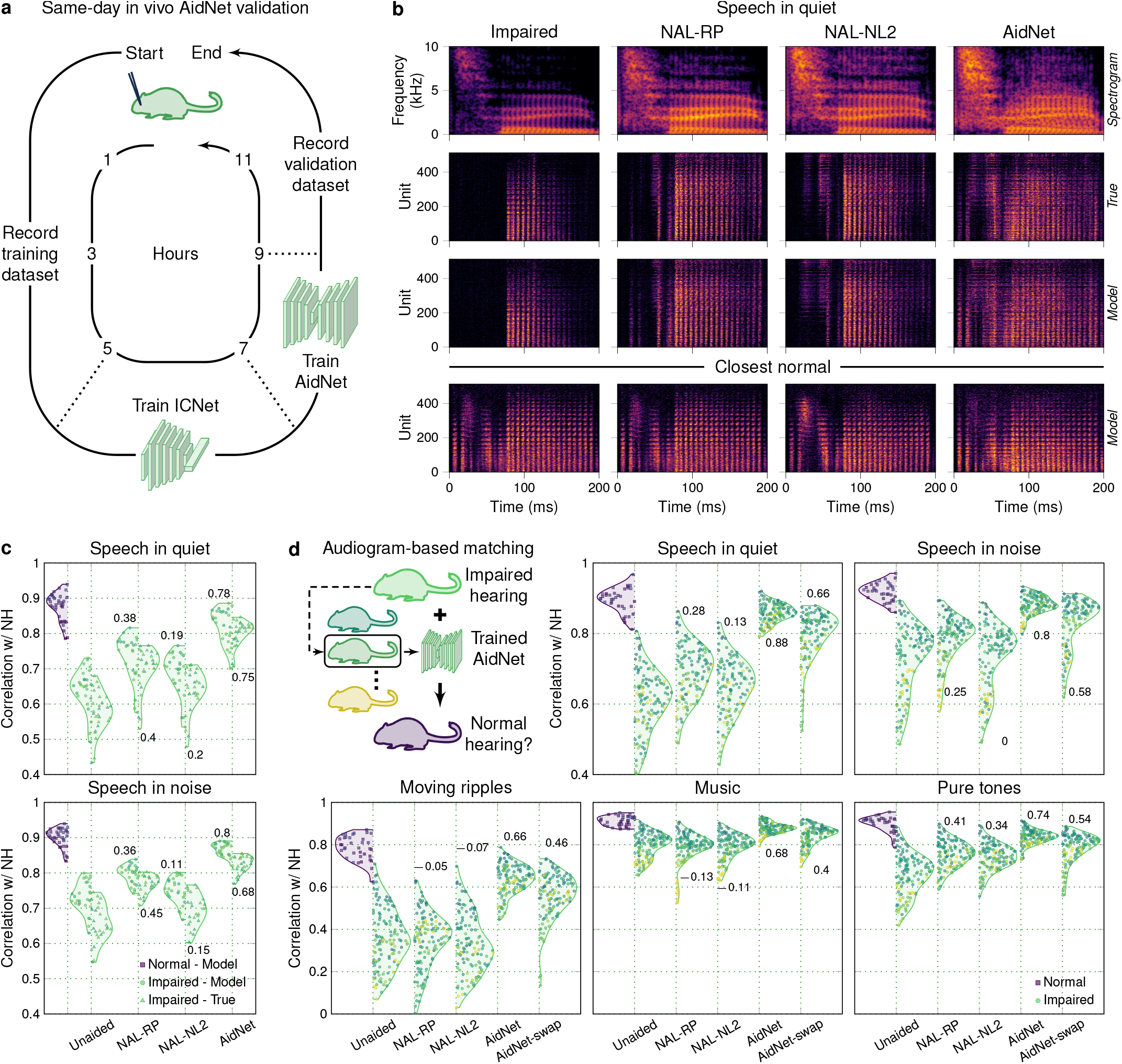
Further validation of AidNet benefit. **a**. Schematic diagram of the experimental protocol that we used to validate an AidNet within a single recording session. **b**. Example spectrograms and responses to speech in quiet at 60 dB SPL. The top row shows sound spectrograms before and after processing. The second and third rows show recorded and simulated HI activity. The bottom row shows the best match (lowest mean absolute error) from the collection of NH responses to each of the HI responses. Neural activity was simulated by sampling from the distribution 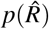 of each ICNet model and was aligned using MCA. Examples for speech in noise are shown in Supplementary Fig. 6. **c**. Restoration results for speech in quiet and in noise from sound corpora that were not part of training. The distributions show the correlation between activity patterns from pairs of 9 NH brains (purple), as well as from pairs of 9 NH and 3 HI brains (yellow and green) for each of the indicated processing conditions. For NH-HI pairs, results are shown for both simulated (circles on left) and recorded (triangles on right) responses. To eliminate neural noise, responses were averaged across repeated presentations (see Methods). Correlation was computed after flattening the responses. The numbers indicate the average restoration index (see Supplementary Table 2). Restoration results for additional sounds are shown in Supplementary Fig. 6. **d**. Simulated restoration results for the full collection of 14 HI animals with and without AidNet swapping. Results are presented as in **c** for five sounds that were not part of training (speech in quiet and in noise, moving ripples, music and pure tones; see Methods). To eliminate neural noise, the expectation over counts of 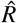 was taken for each ICNet model. The schematic illustrates the pairing of each animal with an AidNet corresponding to one of the other 13 animals with the closest hearing thresholds (AidNet-swap). Results for all pairwise swaps are shown in Supplementary Fig. 7.

Fig. 4b shows sound spectrograms and corresponding real and simulated activity heatmaps in response to speech in quiet for one example animal. As expected, the benchmark amplification strategies restored some of the missing features of the neural responses, but AidNet yielded responses that were closer to normal for both real and ICNet-simulated activity (the bottom row of Fig. 4b shows, for each processing strategy, the closest match to the HI activity from the collection of NH targets). The benefit of AidNet for neural restoration as measured by the correlations between normal and aided activity was consistent with the results observed in silico (Fig. 4c): For recorded activity, the average overall restoration for speech in quiet and in noise was of 0.33, 0.15 and 0.73 for NAL-RP, NAL-NL2 and AidNet, respectively (see Supplementary Table 2). For ICNet-simulated activity for these animals, the results were similar with average overall restoration of 0.29, 0.14 and 0.79.

### AidNets provide generalizable compensation for complex effects of hearing loss

Our approach to AidNet training identifies sound processing that is fully optimized for each animal independently. However, if some of the effects on neural coding are shared across animals with similar hearing loss severity, the processing strategies identified during training may be generalizable to new animals without the need for direct access to the full neural code. To test this, we assessed the restoration achieved when AidNets were swapped across animals with similar hearing thresholds.

For each of our 14 impaired animals, we found the closest match (out of the other 13) based on hearing thresholds and used the second animal’s AidNet when generating responses from the first animal’s ICNet (Fig. 4d). The AidNet advantage was largely retained after swapping (Fig. 4d; top row), with AidNet still outperforming the benchmark strategies by a wide margin (the average overall restoration for speech in quiet and in noise was of 0.27, 0.07, 0.84, and 0.62 for NAL-RP, NAL-NL2, original AidNet, and swapped AidNet, respectively; see Supplementary Table 3 for full statistics and Supplementary Fig. 7 for all pairwise swaps). This result indicates that much of AidNet’s advantage over standard processing strategies comes not from greater individualization, but from its use of fully flexible processing that can correct for complex effects of hearing loss that are common across animals with similar thresholds.

We also tested whether the benefit of AidNet generalized to sounds other than speech and again found that AidNet outperformed the benchmark amplification strategies, both before and after swapping (Fig. 4d; bottom row). For moving ripples, music and pure tones, the average overall restoration was 0.07, 0.05, 0.69, and 0.47 for NAL-RP, NAL-NL2, original AidNet, and swapped AidNet, respectively (see Supplementary Table 2 and Supplementary Fig. 7). This generalized benefit of AidNet was also evident in real activity recorded in vivo (Supplementary Fig. 6). Finally, we sought to ensure that the advantage of AidNet relative to the benchmark strategies was not a result of suboptimal fitting: We refit the benchmark strategies for each animal using different methods for measuring hearing thresholds, but found little effect on neural restoration or hearing sensitivity (Supplementary Fig. 8).

## Discussion

We have demonstrated that closed-loop optimization of hearing aid processing to correct distortions in neural coding is both feasible and effective. We first used large-scale intracranial recordings to train forward models of neural coding (ICNets) to serve as in silico surrogates for normal and impaired brains. We then used the ICNets to train sound processing networks (AidNets) to act as optimal hearing aids by processing sounds as required to elicit normal neural activity in impaired brains. Because AidNets are based on deep neural networks rather than classical signal processing, they can transform sounds in highly non-linear and flexible ways to compensate for the complex effects of hearing loss. One clear example is the injection of low-frequency energy to restore responses to high-frequency sounds (Fig. 3d), which exploits the complexities of cross-frequency cochlear processing. As a result, AidNets outperform standard hearing aid processing (linear amplification and WDRC) by a wide margin in correcting both distorted neural coding and simulated phoneme recognition.

### The advantage of AidNet over standard hearing aid processing

The design of current hearing aids relies on audiometric hearing thresholds for two complementary purposes. The first is to define the focus of their processing, which uses amplification and compression to map the dynamic range of incoming sounds to the reduced dynamic range of a user with hearing loss. The second is to provide a basis for the individualization of their processing for a new user. Because hearing thresholds are a good indication of dynamic range, they can be used to determine the initial settings for the amplification and compression parameters. Our AidNet optimization replaces hearing thresholds with the full neural code for both of these purposes, allowing its processing to compensate for effects of hearing loss beyond just reduced dynamic range and providing additional information for individualization.

Our results show that much of AidNet’s advantage is retained after swapping across animals with similar thresholds, suggesting that thresholds can be an indicator not only of reduced dynamic range, but also of other complex effects of hearing loss. This is not necessarily surprising, as the same cochlear dysfunction that reduces dynamic range is also responsible for many other effects. But it is important because it suggests that while using the full neural code is beneficial for initial development, it may not be necessary for individualization, making the wide use of AidNet-like processing in real devices much more feasible. There are clearly limits to how well hearing aid processing can generalize across individuals (Supplementary Fig. 7), but these limits can be extended by using training datasets that include a wider range of hearing loss as well as different etiologies (e.g., ototoxicity), which may influence the effects on the full neural code that are associated with a given hearing threshold.

It should also be possible to gain additional benefit for new users through individualization based on non-invasive electrophysiology or behavioral feedback (as in the fitting of current hearing aids). If a low-dimensional representation of the full effects of hearing loss can be identified and parameterized, then it can be integrated into AidNet to provide conditional processing and optimized over with limited data (e.g., through Bayesian optimization based on A/B comparisons [25–27]). This approach has already been applied to create forward models that replicate neural coding across a range of hearing impairment [28, 29], as well as to provide sound processing that corrects for a range of audiometric thresholds [30–33].

### Deep learning for closed-loop optimization of sensory stimulation

Our framework to hearing aid design builds on prior approaches that have been developed using cochlear models [10–12]. We extended these approaches to work directly with brain recordings, following work that has used deep learning for neural control in the context of vision. Ponce et al. [13] used a genetic algorithm to search for the latent codes of a pre-trained generative image network which produced images that maximally excited individual neurons in visual cortex. Because this optimization required only a few hundred iterations, it could be performed in real-time while recording from each neuron. Other studies that performed full optimization of raw image pixel values for populations of neurons have used a two-stage approach similar to ours. Bashivan et al. [14] used a pre-trained image classifier as a forward model, choosing the model layer with activations that were most highly correlated with activity in visual cortex to serve as a proxy for the brain. They then used the model to identify images that either maximally excited individual neurons or selectively activated specific neurons within a population. Walker et al. [15] used a similar approach, but trained their own forward model directly on neural recordings.

The use of a forward model makes closed-loop optimization of sensory inputs feasible by allowing stimulus-response iterations to be completed many times faster than in the real brain. And if a high-capacity forward model can be trained directly on neural recordings, this approach allows gaps in the understanding of the underlying system to be overcome with few assumptions. In our case, however, there is an added challenge because the optimization requires aligning activity across different brains without correcting differences related to hearing loss. There are many contexts in which activity must be aligned, but the goal is usually to achieve the best possible match without constraint. For example, for the analysis of neural manifolds or the design of brain-computer interfaces, it is common to learn latent representations that maximize similarity across brains [24] or are invariant to changes across recording sessions [34–36]. To minimize the risk of correcting hearing loss during alignment, we used orthogonal projections to align the output of different ICNets. A more flexible alternative could be to include alignment within the forward models using a framework that enables disentanglement of differences in neural activity that are related to hearing loss from those that are not (analogous to disentanglement of style and position in image generation [37]).

The success of closed-loop neural restoration for hearing aid optimization would pave the way for the extension of the approach to other sensory devices and neurotechnologies, with the most obvious being cochlear implants (CIs). As with hearing aids, CIs have relied on variations of the same stimulation strategy for decades. The performance of CIs in complex listening conditions is often unsatisfactory and strategies such as adding temporal fine structure or activating specific subsets of channels have failed to produce robust results [38, 39]. Assuming an appropriate animal model can be developed, it should be possible to perform closed-loop optimization of CI stimulation, i.e., to identify sound-to-current transformations that are more effective than current stimulation strategies in eliciting the desired downstream neural activity patterns [40]. The translational challenges involved in using this approach for electrical hearing are much more substantial than for acoustic hearing because of the dependence of current spread on cochlear geometry and the neural interface. But with continuing advances in all aspects of CI design [41] and the introduction of other stimulation modalities [42], there may be ways to address these challenges.

## Methods

### Experimental protocol

Experiments were performed on young-adult gerbils of both sexes that were born and raised in standard laboratory conditions. Hearing impairment was induced through exposure to noise at an age of 16-18 weeks. IC recordings were made at an age of 20-24 weeks. All experimental protocols were approved by the UK Home Office (PPL P56840C21).

The study included recordings from 12 animals with normal hearing and 14 animals with hearing impairment. For all of the NH animals and 9 of the HI animals, the full training dataset was recorded. For the other 5 HI animals, only the abridged training dataset was recorded. Two of these animals were used to pilot the in vivo AidNet validation protocol and the other 3 were used for the actual in vivo validation. For the details of the animals used for each of the different analyses in the study, see Supplementary Table 4.

### Noise exposure

Mild-to-moderate sensorineural hearing loss was induced through exposure to high-pass filtered noise with a 3 dB/octave roll-off below 2 kHz at 118 dB SPL for 3 hours [20, 22]. For anesthesia, an initial injection of 0.2 ml per 100 g body weight was given with fentanyl (0.05 mg per ml), medetomidine (1 mg per ml), and midazolam (5 mg per ml) in a ratio of 4:1:10. A supplemental injection of approximately 1/3 of the initial dose was given after 90 minutes. Internal temperature was monitored and maintained at 38.7^◦^ C.

### Preparation for IC recordings

Recordings were made using the same procedures as in previous studies [20–22]. Animals were placed in a sound-attenuated chamber and anesthetized for surgery with an initial injection of 1 ml per 100 g body weight of ketamine (100 mg per ml), xylazine (20 mg per ml), and saline in a ratio of 5:1:19. The same solution was infused continuously during recording at a rate of approximately 2.2 µl per min. Internal temperature was monitored and maintained at 38.7^◦^ C. A small metal rod was mounted on the skull and used to secure the head of the animal in a stereotaxic device. The pinnae were removed and speakers (Etymotic ER-10X) coupled to tubes were inserted into both ear canals. Sounds were low-pass filtered at 12 kHz (except for tones) and presented at 44.1 kHz without any filtering to compensate for speaker properties or ear canal acoustics. Two craniotomies were made along with incisions in the dura mater, and a 256-channel multi-electrode array was inserted into the central nucleus of the IC in each hemisphere.

### Multi-unit activity

Neural activity was recorded at 20 kHz. Multi-unit activity (MUA) was measured from recordings on each channel of the electrode array as follows: (1) a bandpass filter was applied with cutoff frequencies of 700 and 5000 Hz; (2) the standard deviation of the background noise in the bandpass-filtered signal was estimated as the median absolute deviation / 0.6745 (this estimate is more robust to outlier values, e.g., neural spikes, than direct calculation); (3) times at which the bandpass filtered signal made a positive crossing of a threshold of 3.5 standard deviations were identified and grouped to yield spike counts in 1.3 ms bins.

### Auditory brainstem responses

Before beginning the IC recordings, auditory brainstem responses (ABRs) were measured. Subdermal needles were used as electrodes with the active electrodes placed behind the ear over the bulla (one on each side), the reference placed over the nose, and the ground placed in a rear leg. Recordings were bandpass-filtered between 300 and 3000 Hz. The parallel ABR method [43, 44] was used, with randomly timed tones at multiple frequencies presented simultaneously and independently to each ear. The tone frequencies were 1, 2, 4, 8, and 16 kHz. Each tone was five cycles long and multiplied by a Blackman window of the same duration. Tones were presented at a rate of 40 per s per frequency with alternating polarity for 100 s at each intensity. The activity recorded in the 30 ms following each tone was extracted and thresholds for each frequency were defined as the lowest intensity at which the root mean square (RMS) of the median response across presentations was more than twice the RMS of the median activity recorded in the absence of sound. The ABR thresholds for each ear were used to choose animals with symmetric hearing (loss) for this study. Average ABR threshold differences between the two ears of all chosen NH and HI animals were 4.20 ± 2.04 dB and 6.41 ± 1.33 dB, respectively (computed from 1 to 8 kHz).

### Hearing thresholds

As an alternative to ABR-estimated thresholds for fitting the standard hearing aid processing strategies used in this study, thresholds were also estimated from IC activity. The hearing thresholds shown in Fig. 1d (and Supplementary Figs. 5 and 8) were estimated from the neural responses to 50-ms pure tones presented at 7 frequencies (0.25, 0.5, 1, 2, 4 and 8 kHz). Tones were presented at levels from 4 to 85 dB SPL with a step of 9 dB and with 10 ms cosine on and off ramps. Each tone was repeated 8 times in a random order and was followed by 75 ms of silence. The resulting responses were then averaged across repetitions and were used to compute the rate-level functions for each frequency by summing the activity across time bins from 0 to 65 ms and across units. The neural threshold at each frequency was then estimated from the corresponding rate-level curve as follows: (1) a Savitzky-Golay filter of size 3 and order 1 was applied to smooth the rate-level curve; (2) the elbow of the smoothed curve was identified as the first level point at which the 1-sample difference (gradient) of the curve was positive for 3 consecutive levels; (3) baseline activity was computed by averaging the curve values at the 3 levels up to (but not including) the elbow; and (4) the neural threshold was estimated as the level that elicited activity corresponding to 1.25 times the baseline activity. For each individual animal, the ABR thresholds were then used to calibrate the neural thresholds: The difference between the neural and ABR thresholds at 2 kHz (average of the two ears) was computed and applied as an offset value to the neural thresholds across all frequencies.

### Forward models mapping sound to neural activity (ICNets)

DNN models were trained to map a sound waveform *s* to neural activity 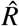 via an inferred conditional distribution of MUA counts 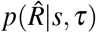 using the convolutional architecture of our previous study [21] that comprised: (1) a SincNet layer [45] with 48 bandpass filters of size 64 and stride of 1, followed by a symmetric logarithmic activation *y* = sgn(*x*) * log(|*x*| + 1); (2) a stack of five 1-D convolutional layers with 128 filters of size 64 and stride of 2, each followed by a PReLU activation; (3) a 1-D bottleneck convolutional layer with 64 filters of size 64 and a stride of 1, followed by a PReLU activation; (4) a cropping layer to eliminate convolutional edge effects as described below; (5) a time module (convolutional layer with 64 filters of size 1, followed by a PReLU activation) that mapped a time input *τ* to the dimensionality of the bottleneck and was multiplied (elementwise) by the cropped bottleneck output; (6) a decoder convolutional layer without bias and *M × N*_*c*_ filters of size 1, where *M* = 512 is the number of neural units and *N*_*c*_ = 5 is the number of classes in the count distribution 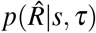; and (7) a softmax activation that produced the probability of each possible count *n* ∈ {0,…, *N*_*c*_ − 1} for each unit in each time bin. All convolutional layers in the encoder included a bias term and used a causal kernel. The additional time input (omitted from the main text for simplicity) was used to account for non-stationarity and had the same sampling rate as the bottleneck output, with each sample defining the relative time at which a corresponding MUA sample was recorded (range of approximately 0-12 hr).

### ICNet training

Models were trained to transform 24,414.0625 Hz sound input frames of 8,192 samples into 762.9395 Hz neural activity frames of 256 samples. Context of 2,048 samples was added on the left side of the sound input frame (total of 10,240 samples) and was cropped after the bottleneck layer (2,048 divided by a decimation factor of 2^5^ resulting in 64 cropped samples at the level of the bottleneck). A crossentropy loss function was used for training: 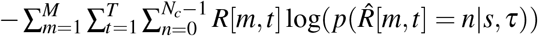, where *T* is total number of time bins and *R* is the recorded neural activity. All neural activity was clipped during training and inference at a maximum value of 4 spikes. Sound inputs were scaled such that an RMS of 0.04 corresponded to a level of 94 dB SPL. Time inputs were expressed in seconds and were scaled by 1/36,000 to match the dynamic range to that of the sound inputs.

The ICNet models were trained on NVidia RTX 4090 GPUs using Python and Tensorflow [46]. A batch size of 50 was used with the Adam optimizer and a starting learning rate of 0.0004. All trainings were performed so that the learning rate was halved if the loss in the validation set did not decrease for 2 consecutive epochs. Early stopping was used and determined the total number of training epochs if the validation loss did not decrease for 4 consecutive epochs. To speed up training, the weights from the encoder of a previous version of ICNet [21] were used to initialize the models. This resulted in an average of 20.1 ± 6.4 epochs for the 23 trained ICNet models used in Figs. 1-3 (see Supplementary Table 4).

### ICNet training dataset

The sounds that were used for training the ICNet models are described in detail in our previous study [21]. The training dataset totaled to 7.83 hr and included a mixture of speech in quiet and in noise, processed speech, music and moving ripples. The order of the sound presentation was randomized for each animal during recording, with 10% of the sounds randomly chosen to form the validation set during training.

### Abridged training dataset

A subset of all sounds was used to train ICNet models for the animals used for in vivo validation (Fig. 4) and for two of the remaining HI animals (see Supplementary Table 4). The abridged training dataset was generated by keeping only the first 180 s of each speech segment (60% of total duration), while preserving the total duration of the remaining sounds. The resulting dataset totaled to 5.78 hr and included 2.95 hr of (processed and unprocessed) speech and 2.83 hr of music and ripples. As before, 10% of the sounds were randomly chosen to form the validation set during training.

### ICNet evaluation

To simulate neural activity using the trained ICNet models, the decoder parameters were used to define a categorical probability distribution with *N*_*c*_ classes using the Tensorflow Probability toolbox [47]. The distribution was then sampled from to yield simulated neural activity across time bins and units for the analyses shown in Fig. 1. The time input to the ICNet models was matched to the time that the responses to each evaluation sound were recorded for each animal.

### ICNet evaluation dataset

The sounds that were used for evaluating the ICNet models were not part of the training dataset and are described in detail in our previous study [21]. Each sound segment was 30 s in duration and was presented twice in successive trials. The sounds were: (1) a speech segment from the UCL SCRIBE dataset consisting of sentences spoken by a male talker presented at 60 dB SPL; (2) a speech segment from the UCL SCRIBE dataset consisting of sentences spoken by a female talker presented at 85 dB SPL in hallway noise from the Microsoft Scalable Noisy Speech dataset at 0 dB SNR; (3) dynamic moving ripples with frequencies between 4.7 and 10.8 kHz presented at 85 dB SPL; and (4) three seconds from each of 10 mixed pop songs presented at 75 dB SPL.

### Predictive power metrics

To assess the predictive power of the ICNet models (Fig. 1c), we computed a metric using successive trials (to minimize any impact of non-stationarity) for the 20 animals for which this was possible (10 NH and 10 HI; see Supplementary Table 4). The metric is formulated as the percentage of explainable correlation that the model explained across all units and was computed as follows:

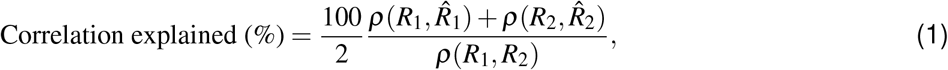

where *R*_1_, *R*_2_ ∈ ℕ^*M×T*^ are the recorded neural activity for each of the two trials, 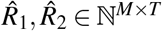 are the predicted neural activity, and is the correlation coefficient. The responses were flattened (collapsed across time and units into one dimension) before the correlation was computed.

### Neural alignment

Four methods were used to align neural activity from different brains. The alignment was performed on the recorded responses to a subset of the ICNet abridged training dataset. For each sound of the dataset, the responses between 16.77 and 83.88 s after sound onset (200 frames of 256 samples starting from the 50th frame) were used to minimize the effect of onset dynamics. One NH brain was chosen as the reference for all comparisons, with the remaining brains aligned to this reference. Given the response matrices *R*_*i*_, *R*_*re f*_ ∈ ℕ^*M×T*^ for each brain and the reference brain, respectively, a transformation was sought to make the aligned response *R*_*i*→*re f*_ as close as possible to *R*_*re f*_, subject to constraints that varied across methods.

### Linear Regression

Least squares optimization from the Procrustes library in Python [48] was used to obtain a transformation matrix *T*_*i*_ ∈ ℝ^*M×M*^ that minimized ||*R*_*i*_*T*_*i*_ −*R*_*re f*_ ||^2^ without constraint, where || · ||_*F*_ is the Frobenius norm.

### Procrustes

Singular value decomposition was used to obtain a transformation matrix *T*_*i*_ ∈ ℝ^*M×M*^ that minimized 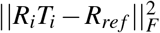 subject to 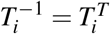 and |*T*_*i*_| = 1.

### Optimal Transport

The earth mover’s distance algorithm from the Optimal Transport toolbox in Python [49] was used to derive a permutation matrix *T*_*i*_ (i.e., the optimal channel reordering of *R*_*i*_) that minimized the Wasserstein distance between the responses *R*_*i*_*T*_*i*_ and *R*_*re f*_.

### Maximum Covariance Analysis

Singular value decomposition was used to define the vectors *U*_*i*_, *V*_*i*_ that factorize the covariance matrix 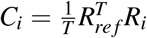 as 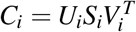, where *S*_*i*_ is a diagonal matrix and *U*_*i*_, *V*_*i*_ are orthonormal unitary vectors. The aligned responses were then computed as *R*_*i*→*re f*_ = (*D*_*i*_(*R*_*i*_*V*_*i*_)^*T*^)^*T*^, where 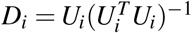.

### Alignment evaluation

Speech from the test subset of the TIMIT corpus [50] was used to evaluate the alignment methods (Fig. 2 and Supplementary Figs. 1-2). The speech segment was 35 s in duration and was presented: (1) at 50 dB SPL in quiet; and (2) at 65 dB SPL mixed at 0 dB SNR with the restaurant noise from the Microsoft Scalable Noisy Speech Dataset [51]. The sounds were segmented into frames of 65,536 samples with context of 4,096 samples on both sides and were given as inputs to the ICNet models to simulate neural activity (MUA frames of 2,048 samples after cropping the added context). A time input corresponding to a random number between 1 and 2 hours was used for the ICNet models to simulate responses at the beginning of each recording and eliminate any non-stationarity effects [21]. The simulated MUA responses shown in Fig. 2 are the expectation over counts 𝔼_*c*_ of 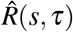 after alignment with each of the methods explained above.

### Hearing aid processing models mapping sound to sound (AidNets)

The hearing aid model, AidNet, comprised a symmetric encoder-decoder DNN architecture based on the Wave-U-Net model [52] that was trained to process sound end-to-end with 18 convolutional layers in total. The encoder consisted of 9 1-D convolutional layers that had a kernel size of 7, stride of 1, and ascending number of filters: 32, 32, 64, 64, 64, 128, 128, 128, and 256 filters, respectively. Each convolutional layer was followed by a PReLU activation and a decimation layer that kept every second temporal sample and downsampled the signal by a factor of 2. The decoder mirrored the layers of the encoder to upsample the output of the last encoder layer back to the original dimensionality of the sound. It consisted of upsampling layers that applied bilinear interpolation to double the temporal resolution, and skip connections (linking each encoder layer to the respective decoder layer), followed by convolutional layers and PReLU activations. All convolutional layers included a bias term and used a non-causal kernel, with the input to each convolutional layer padded with zeros at the beginning and end to maintain the size of the temporal dimension.

### AidNet training

Fig. 3a illustrates the framework that was used to train an AidNet model for a given HI brain. The sound input *s* was given to a NH ICNet to produce a NH response 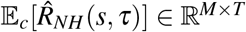, where *M* is the number of units, *T* is the number of time bins, and 𝔼_*c*_ denotes expectation over counts. For all AidNet training, the time input *τ* for the ICNet models was set to a random number between 1 and 2 hours in each batch and is omitted from here on for clarity. Nine NH ICNets were used as the target during training, with one randomly chosen to provide the model weights in each training batch. Only three of these nine ICNets were used in the validation stage of the training, with one of the three randomly chosen before each validation batch.

The sound input *s* was also given to AidNet to produce a processed sound output *s*^*^, subsequently used as input to a HI ICNet to produce a response 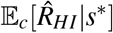. The context of the original sound was always used in place of the AidNet-processed context to ensure that neural restoration could be achieved without manipulation of the context. The two responses 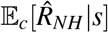 and 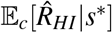 were aligned to the reference brain using MCA, resulting in the final responses 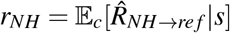 and 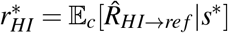 The weights of the AidNet model were then updated via backpropagation based on the differences in these two responses as defined by a composite loss function (see below). The responses were also given as input to an automatic speech recognition (ASR) backend to additionally target the restoration of simulated phoneme recognition, and training was done both with and without this ASR backend to assess its impact. The ICNet and ASR backend models were frozen during AidNet training.

The AidNet models were trained on NVidia RTX 4090 GPUs using Python and Tensorflow [46]. A batch size of 30 was used with the Adam optimizer and a starting learning rate of 0.0004. All trainings were performed so that the learning rate was halved if the loss in the validation set did not decrease for 3 consecutive epochs. Early stopping was used if the validation loss did not decrease for 9 consecutive epochs. This resulted in an average of 22.1 ± 3.1 epochs for the 11 AidNet models of Fig. 3 (21.5 ± 3.4 for the 11 AidNet models of Supplementary Fig. 4 trained without the ASR backend). To speed up training, an AidNet model was trained from scratch for one randomly chosen HI animal and the resulting weights were used to initialize all AidNet models.

#### AidNet training dataset

The dataset that was used to train AidNets was mainly composed of speech, but also contained music and moving ripples, with all sounds resampled to 24,414.0625 Hz and calibrated to randomly selected levels between 40 and 90 dB SPL in steps of 10 dB. Sound inputs were scaled such that an RMS of 0.04 corresponded to a level of 94 dB SPL. For speech, the train subset of the TIMIT corpus [50] and the dev-clean subset of the LibriTTS corpus [53] were used, resulting in 4.12 hr and 6.35 hr of data, respectively. Sentences were randomly mixed with the 18 noise types from the DEMAND dataset [54] at SNRs of -6, -3, 0, 3, 6, 9, 12, 15, and 100 dB. Only positive SNRs were used for levels below 70 dB SPL to construct a more naturalistic dataset.

For music, 10 full pieces of rock and classical music were included with a total duration of 0.58 hr. Pop songs taken from the musdb18 dataset [55] were also used in full mixed form, as well as in stem form with isolated tracks for drums, bass, vocals and other (e.g., guitar, keyboard) for a total duration of 2.34 hr. Band-limited dynamic moving ripple sounds (created as described in [21]) were also included. The lowest frequency was either 4.7 kHz or 6.2 kHz and the highest frequency was always 10.8 kHz. The total duration of the ripples in the dataset was 0.67 hr.

All sounds were segmented to sound input frames of 65,536 samples (without overlap) to form the training dataset. Context of 4,096 samples was added on both sides of the sound input frames. The phonemic transcriptions of the speech corpora were downsampled to the sampling frequency of the neural activity (762.9395 Hz) and were grouped into 40 classes as specified below (15 vowels, 24 consonants and a silence class). Phonemic transcriptions for the LibriTTS corpus were obtained using the Montreal Forced Aligner from https://github.com/kan-bayashi/LibriTTSLabel.

#### Phoneme analysis

The phonemic transcriptions of the TIMIT dataset were grouped into 40 classes that comprised 15 vowels, 24 consonants and a silence class that included all pauses and the glottal stops in the dataset [56]. These classes were then grouped into two categories for analyzing the results: (1) vowels and vowel-like consonants (/a/, /xq/, /xa/, /c/, /xw/, /xy/, /xr/, /xe/, /e/, /xi/, /i/, /o/, /xo/, /xu/, /u/, /m/, /n/, /xt/, /r/, /l/, /w/, /y/); and (2) remaining consonants (/b/, /xc/, /d/, /xd/, /f/, /g/, /h/, /xj/, /k/, /xg/, /p/, /s/, /xs/, /t/, /v/, /z/, /xz/). Alignment was evaluated separately on the segments of the MUA responses that corresponded to the phonemes in the two categories. Phoneme transcriptions were downsampled to the MUA sampling frequency and were shifted by 6.5 ms to account for the overall latency in IC activity.

### AidNet loss function

AidNets were trained to identify the sound processing that minimized a composite loss function. The loss function included three terms that were computed in each training step and reflected differences between: (1) impaired and normal neural responses; (2) unprocessed and AidNet processed sound; and (3) impaired and normal phoneme recognition. All loss components were computed after cropping the added context.

#### Neural restoration

For each batch, a loss term *L*_*r*_ was computed between the responses of a given HI ICNet 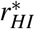 and one of the nine NH ICNets *r*_*NH*_. The loss term combined several metrics to assess different aspects of the differences between the normal and impaired activity:

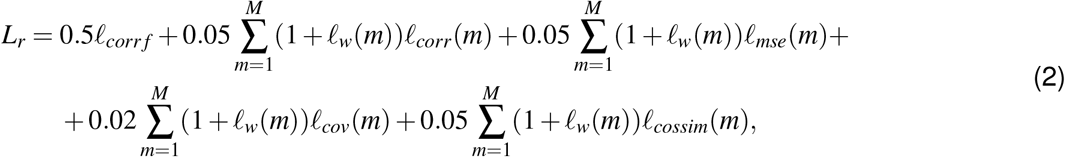

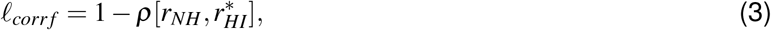

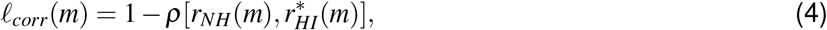

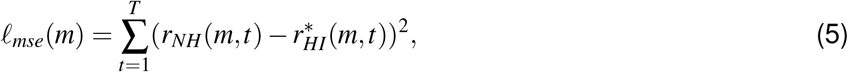

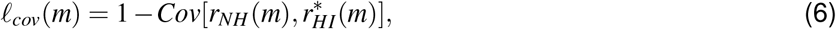

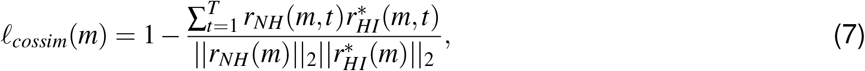

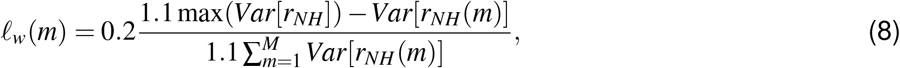

where *M* is the number of units, *T* is the number of time bins, *ρ* is the correlation coefficient, and *Var* and *Cov* are the variance and covariance computed across time bins. The responses were flattened (collapsed across time and units into one dimension) to compute *ℓ*_*corr f*_. Channel-wise metrics were weighted by *ℓ*_*w*_, a weight computed from the overall variance of each channel to ensure that all channels contributed appropriately to the overall loss.

#### Sound processing

An additional loss term *L*_*s*_ was included to minimize differences between the unprocessed and processed sound:

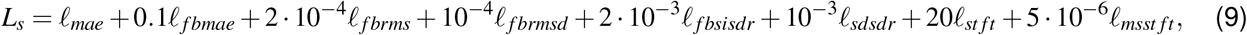

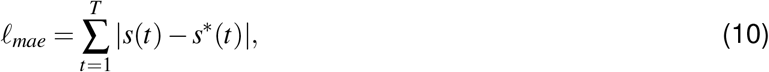

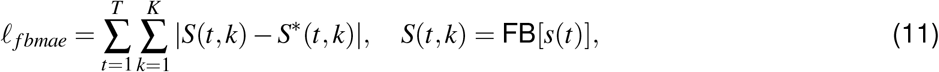

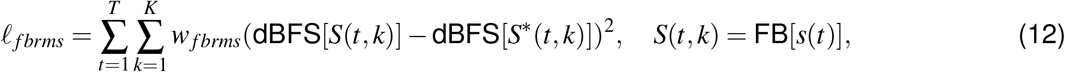

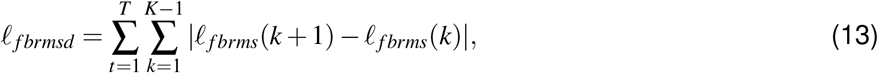

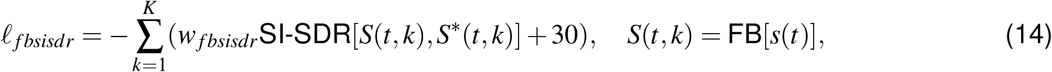

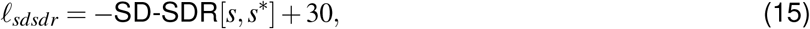

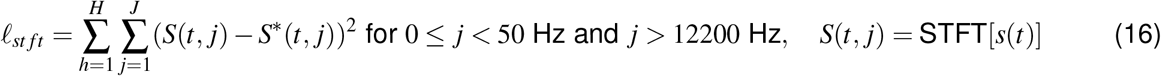

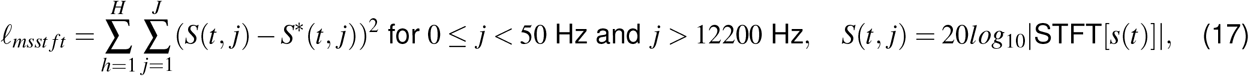

where FB is a filterbank with *K* = 64 SincNet filters [45] spaced logarithmically between 80 and 12000 Hz and 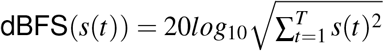. The Short-Time Fourier transform (STFT) used a temporal window of 64 samples, a step size of *H* = 32 samples, and a fast Fourier transform (FFT) size of *J* = 512 samples. SI-SDR and SD-SDR correspond to the scale-invariant and scale-dependent signal-to-distortion ratios and were computed across the time dimension based on equations 3-8 in [57]. For the loss terms of Eqs. *12* and *14*, a sigmoidal weighting factor was applied across the 64 frequency bands to proritize the processing of low frequencies:

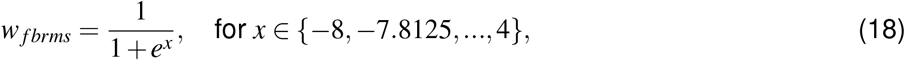

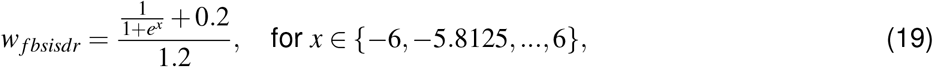

All weights were chosen to ensure a lower overall contribution of *L*_*s*_ relative to *L*_*r*_.

#### Phoneme recognition

A final optional loss term *L*_*p*_ was computed using a pretrained ASR model (described below) to prioritize the restoration of neural features that are relevant for speech perception. The normal and impaired responses (*r*_*NH*_ and 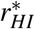) were used as inputs to the ASR backend in each training step to simulate normal and impaired phoneme recognition, respectively (output probabilities of 40 phoneme classes across time [56]). *L*_*p*_ was then computed as the crossentropy between these two outputs and scaled by 0.1 so that its contribution to the overall loss was similar to that of *L*_*s*_. The *L*_*p*_ was computed only on segments containing speech, thus omitting silent parts of speech as well as other sounds.

### ASR model training

The ASR model was trained to predict phonemes from MUA responses using a DNN architecture based on Conv-TasNet [58] that comprised: (1) a convolutional layer with 64 filters of size 3 and no bias, followed by a PReLU activation; (2) a group normalization layer that divided the channels into 32 groups and normalized each by their overall mean and variance, followed by a convolutional layer with 128 filters of size 1; (3) a block of 8 dilated convolutional layers (dilation from 1 to 2^7^) with 128 filters of size 3, including PReLU activations and residual skip connections in between; (4) a convolutional layer with 256 filters of size 1, followed by a sigmoid activation; (5) an output convolutional layer with 40 filters of size 3, followed by a softmax activation. The final output of the ASR model yielded the probabilities of 40 phonemes classes across each MUA time bin (after cropping the added context from both sides). All convolutional layers were 1-D and used a non-causal kernel with a stride of 1. Inputs to each convolutional layer were padded symmetrically with zeros to maintain the size of the temporal dimension.

The training dataset totalled to 57.13 hr and included the train subset of the TIMIT corpus [50] and the train-100, dev-clean and dev-other subsets of the LibriTTS corpus [53]. Sentences were resampled to 24414.0625 Hz, calibrated to randomly selected levels between 30 and 100 dB SPL in steps of 5 dB, and randomly mixed with the 18 noise types from the DEMAND dataset [54] at SNRs of -30, -20, -10, 0, 10, 20, 30, and 100 dB. The phonemic transcriptions of the two corpora were downsampled to the sampling frequency of the neural activity (762.9395 Hz) and were grouped into 40 classes (as explained above). Sentences were then segmented into sound input frames of 73,728 samples to form the training dataset (windows of 65,536 samples with context of 4,096 samples on both sides).

The ASR model was trained to recognize phonemes from the responses of 12 NH and 9 HI ICNets (smaller points in the last row of Supplementary Fig. 4). In each training step, the sound input frames were given to a randomly chosen ICNet to generate activity across 512 neural units at 762.9395 Hz (output frames of 2,048 samples with context of 128 samples on both sides). The neural response was aligned using MCA (as described above) and was given to the ASR backend to predict the probabilities of the 40 phoneme classes across time (output frames of 2,048 samples at 762.9395 Hz after cropping the context). The ASR model was trained to minimize the crossentropy loss between the probability outputs and the phoneme transcriptions in the datasets. The loss was computed only on the segments corresponding to speech (discarding silent segments).

### In silico evaluation

Neural restoration after hearing aid processing was first assessed using the same speech sounds as described above for the alignment comparisons (Fig. 3). The original sounds were given as inputs to all NH and HI ICNets to obtain the responses for speech in quiet and in noise. The same sounds were then given as inputs to each AidNet model followed by the corresponding HI ICNet model to yield the impaired responses after AidNet processing. The expectation over counts 𝔼_*c*_ of 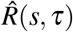 was taken and all responses were aligned using MCA. A time input corresponding to a random number between 1 and 2 hours was used for the ICNet models.

#### Benchmark amplification strategies

Two standard amplification strategies were used as benchmarks for comparison against the restoration achieved by AidNet: (1) a linear strategy (NAL-RP) that applied a frequency-dependent amplification [59]; and (2) a wide dynamic range compression (WDRC) strategy (NAL-NL2) that applied a frequency- and level-dependent amplification [60]. The implementations of the two strategies from the openMHA toolbox in MATLAB were used [61]. Hearing thresholds estimated from the neural responses to pure tones were used to fit the hearing aid strategies for both ears. All sounds were upsampled to 44,100 Hz, and rescaled such that an RMS of 1 corresponded to a level of 110 dB SPL. Sound inputs were then analyzed using a Short-Time Fourier Transform with a window size of 128 samples and 50% overlap, and the resulting frequency bins were grouped into 18 bands spaced logarithmically on a Bark scale. Time constants of 20 ms and 100 ms were used for attack and release, respectively. The time constants were chosen from among a number of tested combinations commonly used in hearing aids because they yielded the best restoration performance in initial tests.

### In vivo evaluation

For the in vivo validation of AidNet, an experimental protocol was designed that combined the collection of neural recordings and the training of the deep learning models within the same day (Fig. 4a). The experimental protocol lasted approximately 12 hours and comprised the following steps:

1. Hours 0-5: Recording neural responses from a HI animal to the abridged ICNet training dataset
2. Hours 5-6: Training an ICNet model using the recorded neural data
3. Hours 6-8: Training an AidNet model to restore the impaired neural activity simulated by the trained ICNet model
4. Hours 7-8: Recording neural responses to the evaluation dataset (original sounds and sounds processed with the benchmark amplification strategies using different fitting procedures)
5. Hours 8-9: Recording neural responses to the evaluation dataset (sounds processed by AidNet)
6. Hours 9-12: Recording additional neural responses for all evaluation sounds and processing conditions

#### AidNet evaluation dataset

The evaluation dataset used in Fig. 4 comprised the four main evaluation sounds described above for ICNet evaluation (speech in quiet, speech in noise, moving ripples and music) presented at 60, 70, 85 and 75 dB SPL, respectively, as well as a different set of the pure tone stimuli that were used to define hearing thresholds (50-ms pure tones with frequencies from 0.13 to 10.81 kHz with a step of 0.2 octaves, presented at levels between 4 and 76 dB SPL with a step of 9 dB). Multiple trials of the five evaluation sounds were presented to three noise-exposed animals, with 20 trials for the speech sounds, 10 trials for ripples and music, and 8 trials for pure tones. The same sounds were then given as inputs to the corresponding ICNet model of each animal to simulate neural activity by sampling from the output distribution of each model. A time input corresponding to a random number between 5 and 6 hours was used for the models to match as much as possible the recorded time of the real responses without exceeding the times used in training. Correlation was computed on the (aligned) responses after averaging across all available trials (Fig. 4 and Supplementary Fig. 6). For the pure tones, the first 52.4 ms of the responses to all tonal stimuli were used. These five sounds were also used to simulate the neural activity of all HI animals for Fig. 4d, using the respective ICNet models and time inputs corresponding to random numbers between 1 and 2 hours.

#### Alternative benchmark amplification fitting

During the in vivo evaluation, the two benchmark hearing aid strategies were fit using several different threshold measurements. For the results shown in Fig. 4, the average neural thresholds across both ears were used (as for the in silico evaluation). Results for alternatives (including the neural threshold for each ear, the average ABR thresholds across both ears, and the ABR threshold for each ear) are shown in Supplementary Fig. 8.

### Statistics

Confidence intervals for all reported values were computed using 1000 bootstrap samples. For assessing model performance (explainable correlation explained) in Fig. 1c, bootstrapping was performed across units for each ICNet. For the restoration performance (Supplementary Tables 1-3), bootstrapping was performed across individual HI animals as follows: (1) the average restoration index for each individual HI animal was computed from the correlation or phoneme recognition results by averaging across NH animals; (2) bootstrap sampling was then performed over HI animals and the 2.5%, 50%, and 97.5% percentiles over the samples were computed (corresponding to the reported median and 95% confidence intervals). The restoration performance reported in Figs. 3-4 and Supplementary Figs. 6-8 corresponds to the average of the restoration index values in step 1 (before bootstrapping). Bonferroni correction was applied to the *p* values of Supplementary Tables 1-3 based on the number of comparisons for each sound.

## Data availability

The four main sounds that were used to evaluate the ICNet and AidNet models (speech in quiet, speech in noise, moving ripples, music) are available via https://doi.org/10.5281/zenodo.19924033. The four sounds are provided together with the corresponding ICNet responses of Fig. 1, as well as with the ICNet responses of Fig. 4 before and after processing with all sound processing strategies. The dataset can be used to replicate the results of Fig. 1c, Fig. 4c-d and Supplementary Fig. 6b. Researchers seeking access to the full set of neural recordings for research purposes should contact the corresponding author via e-mail to set up a material transfer agreement.

## Code availability

The AidNet model for the example HI animal used throughout this work (dotted green line in Fig. 1d; middle column in Supplementary Fig. 5) is available via https://doi.org/10.5281/zenodo.18407090 or https://github.com/fotisdr/AidNet_example. A Jupyter notebook is included with a simple usage example for the AidNet model and sound examples. A non-commercial, academic UCL license applies.

## Acknowledgments

The authors thank Maneesh Sahani, Fan-Gang Zeng, Mohsen Imani, Marius Pachitariu and Tobias Goehring for their advice. This work was supported by UK MRC UKRI3206, EPSRC EP/W004275/1, BBSRC BB/Y008758/1 and MRC MR/W019787/1.

## Author contributions

**F.D**.: Conceptualization, Data Curation, Formal analysis, Investigation, Methodology, Software, Validation, Visualization, Writing: Original Draft Preparation; **L.P**.: Conceptualization, Methodology, Software; **S.S**.: Investigation; **Y.X**.: Investigation; **T.G**.: Investigation; **A.F**.: Conceptualization, Project administration, Software, Supervision; **N.A.L**.: Conceptualization, Data Curation, Funding acquisition, Investigation, Methodology, Project Administration, Resources, Supervision, Writing - Review & Editing

## Competing interests

N. A. L. is a co-founder of Perceptual Technologies. A patent application (GB2412076.8) has been filed by UCL Business (UCLB) on the basis of the research presented in this manuscript.

---

**Supplementary Fig. 1.**
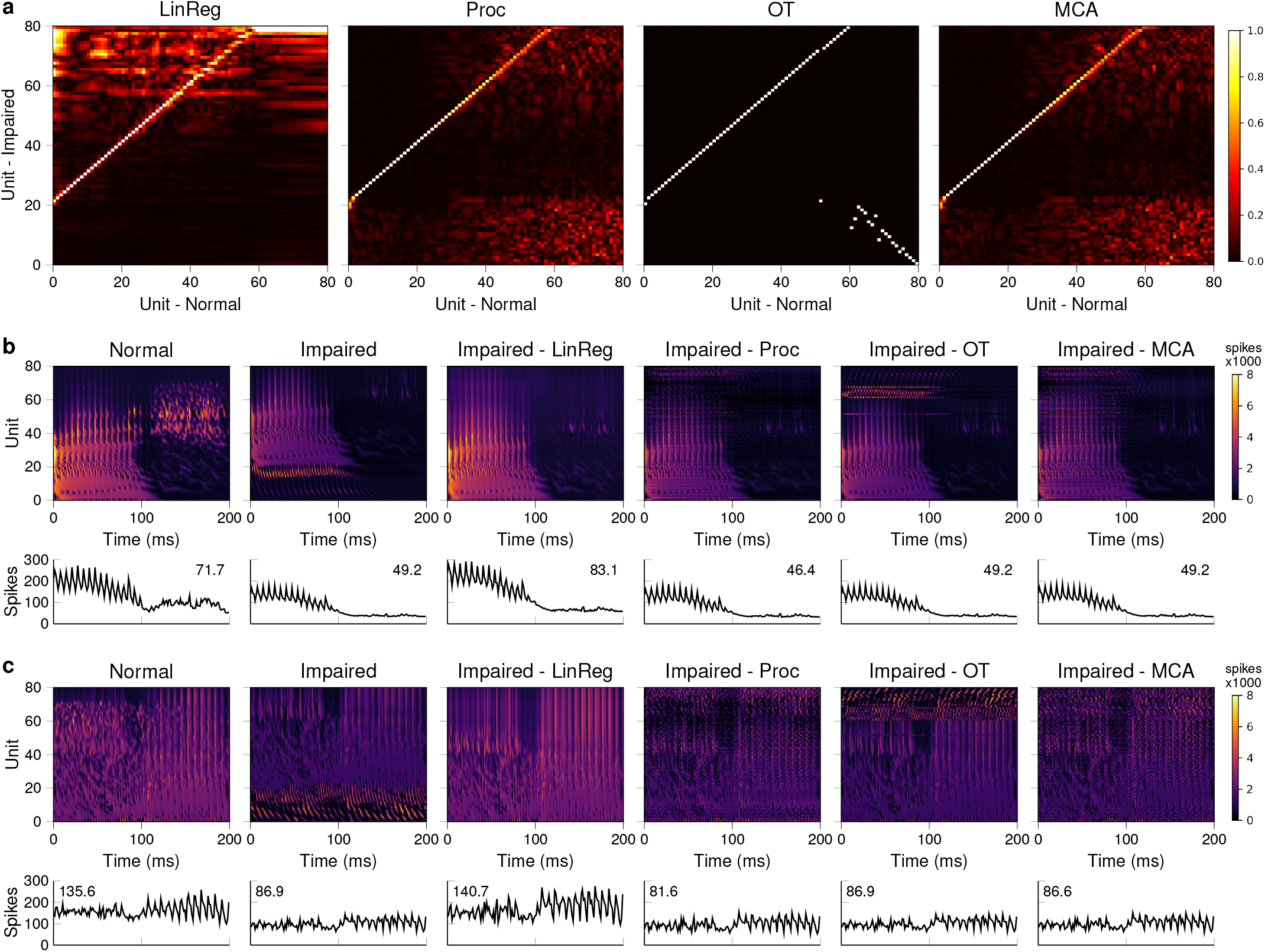
Aligning simulated auditory nerve responses. The ICNet abridged training dataset was used as input to an analytical cochlear model [62, 63] to simulate normal-hearing (NH) and hearing-impaired (HI) responses of the auditory nerve. Normal responses were simulated by summing the firing rates of 13 high-, 3 mid- and 3 low-spontaneous-rate auditory-nerve fibers for each frequency channel. To simulate the impaired responses, a sloping high-frequency loss (starting at 1 kHz and reaching a 35 dB loss at 8 kHz) was imposed at the cochlear stage, and the firing rates of 10 high-spontaneous-rate auditory-nerve fibers were summed (corresponding to a complete loss of mid- and low-spontaneous-rate fibers and a loss of 23% of high-spontaneous-rate fibers). Misalignment of the simulated responses was then introduced by selecting subsets of 80 from the total number of simulated channels (201; CFs ranging from 0.1 to 12 kHz with logarithmic spacing), starting either from the beginning (0:2:160) or the end (41:2:201) to choose non-overlapping channels with slightly different frequency ranges. The simulated responses were downsampled to match the time bin size of our neural responses and alignment was performed in the same way as for Fig. 2 (see Methods). **a**. Transformation matrices derived for the 4 different alignment methods applied to align the HI responses to the NH responses. For MCA, the transformation matrix was defined as the product of the two projection matrices *V*_*i*_*D*^*T*^ (see Methods). To aid visualization, the absolute value of all transformation matrices are shown. All alignment methods do well in matching up units from the CF region that was represented in both responses (units 1-60 for normal and 21-80 for impaired) but differ in how they map between the CF regions that were not present in both responses. Panels **b-c** show example responses with the different alignments for the speech segment of Fig. 2 presented in quiet and in noise. The bottom row shows activity summed across units, with the numbers indicating the total activity (in x1000 spikes) averaged across all time bins for the entire sound. Linear regression (‘LinReg’) is able to correct some of the effects of hearing loss, making it unsuitable for our purposes. Optimal transport (‘OT’), which is constrained to permutations, is forced to match low-CF channels from the impaired response with high-CF channels from the normal response, which is not ideal. Procrustes (‘Proc’) and MCA deal with the misalignment more appropriately by smoothing out responses from the CF regions that were not represented in both responses, with MCA resulting in only negligible differences in summed activity.

**Supplementary Fig. 2.**
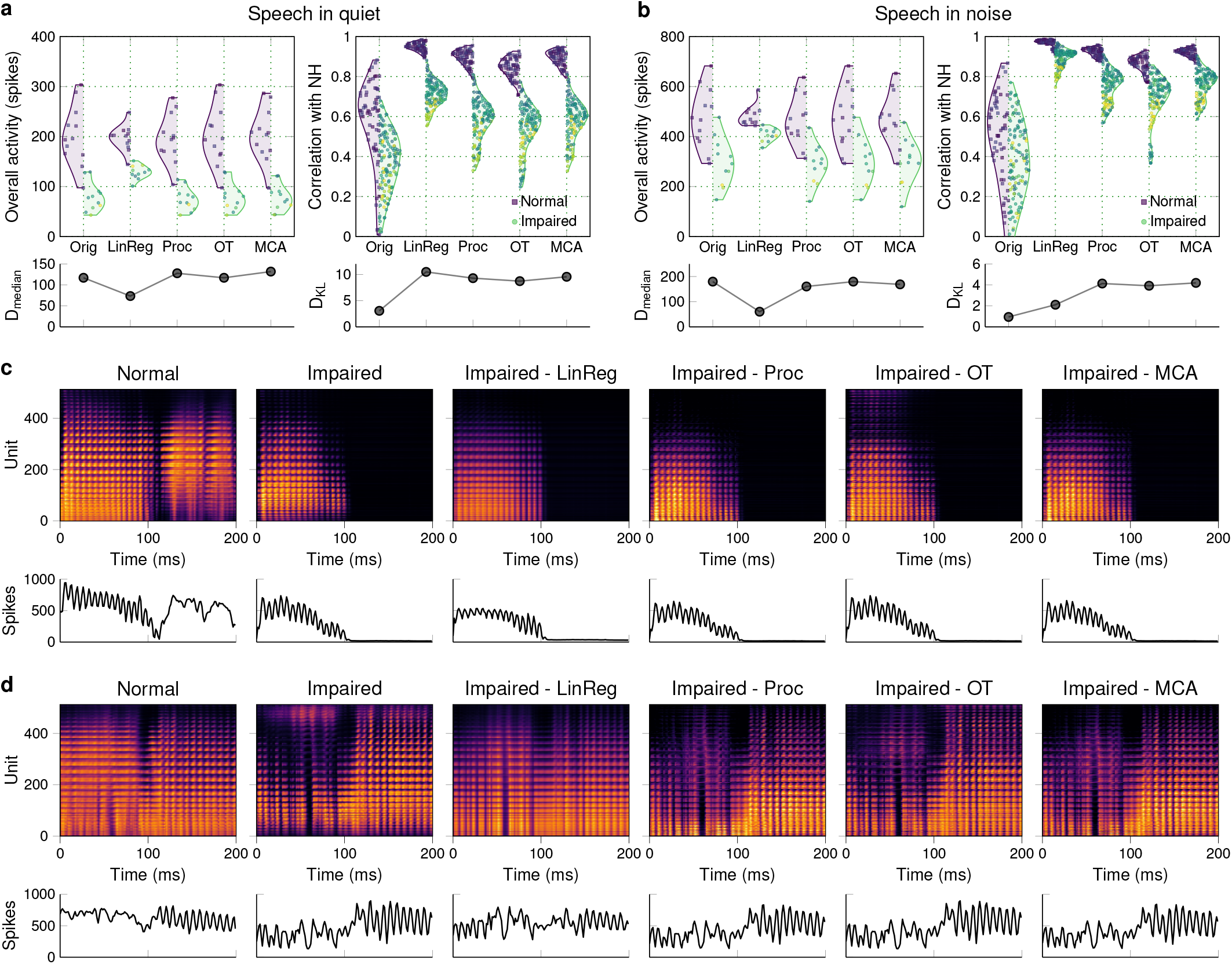
Aligning neural recordings with different methods. **a**. The results of Fig. 2b for speech in quiet at 50 dB SPL are presented before (Orig) and after alignment with all tested methods: Linear Regression (LinReg), Orthogonal-Rotational Procrustes (Proc), Optimal Transport (OT), and Maximum Covariance Analysis (MCA). **b**. Results for speech at 65 dB SPL mixed with restaurant noise at 0 dB SNR, presented as in Fig. 2c. **c**. Example responses to speech in quiet presented as in Fig. 2d. **d**. Example responses to speech in noise presented as in Fig. 2e.

**Supplementary Fig. 3.**
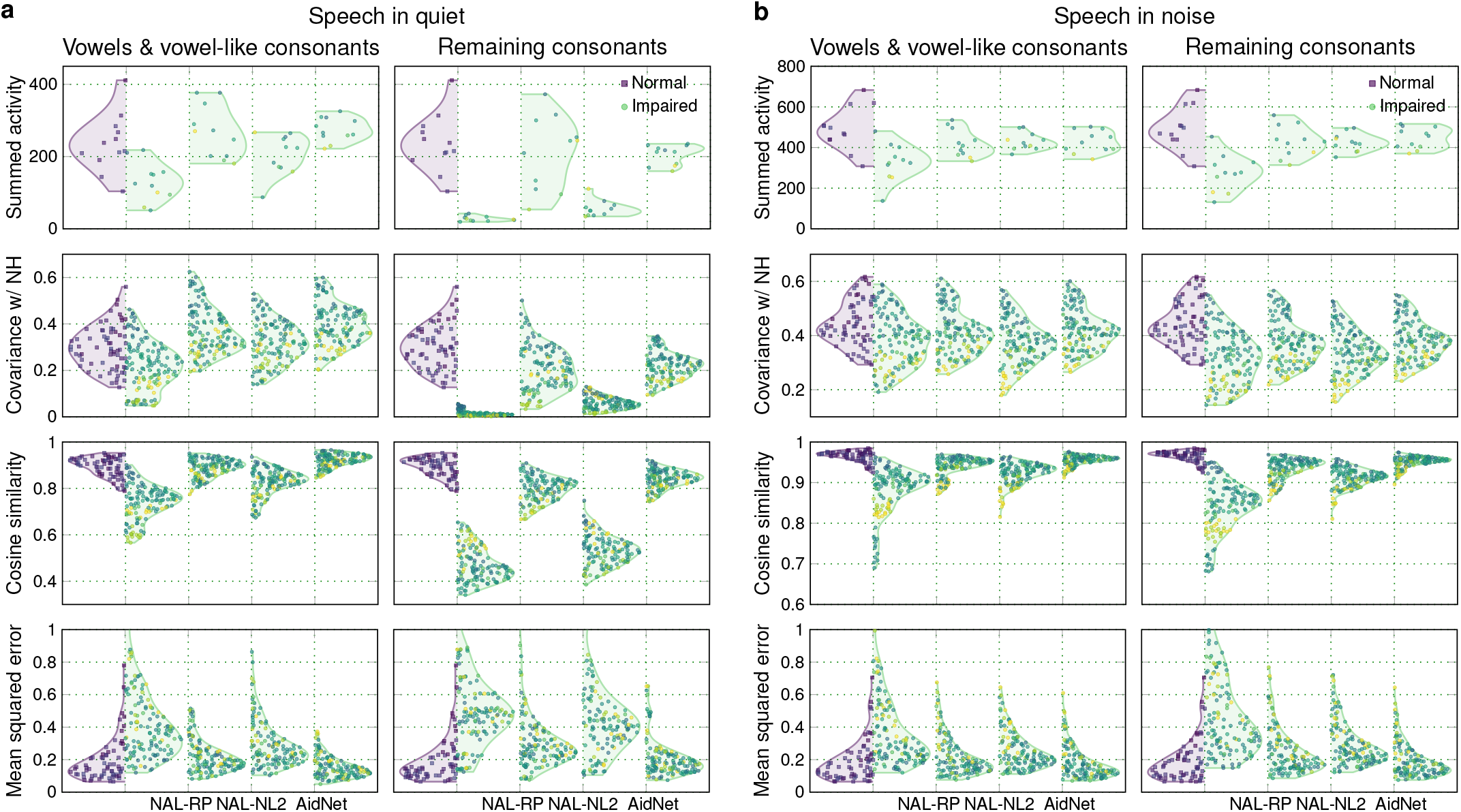
Comparison of different neural metrics. Results for all the similarity metrics that were used in AidNet training are shown for all sound processing strategies (see Methods for metric definitions). The expectation over counts of 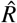 was taken for each ICNet model, and the metrics were computed after applying MCA alignment and flattening the responses. **a**. Results for the vowel and consonant parts of speech presented at 50 dB SPL in quiet. **b**. Results for speech mixed with restaurant noise at 0 dB SNR and presented at 65 dB SPL.

**Supplementary Fig. 4.**
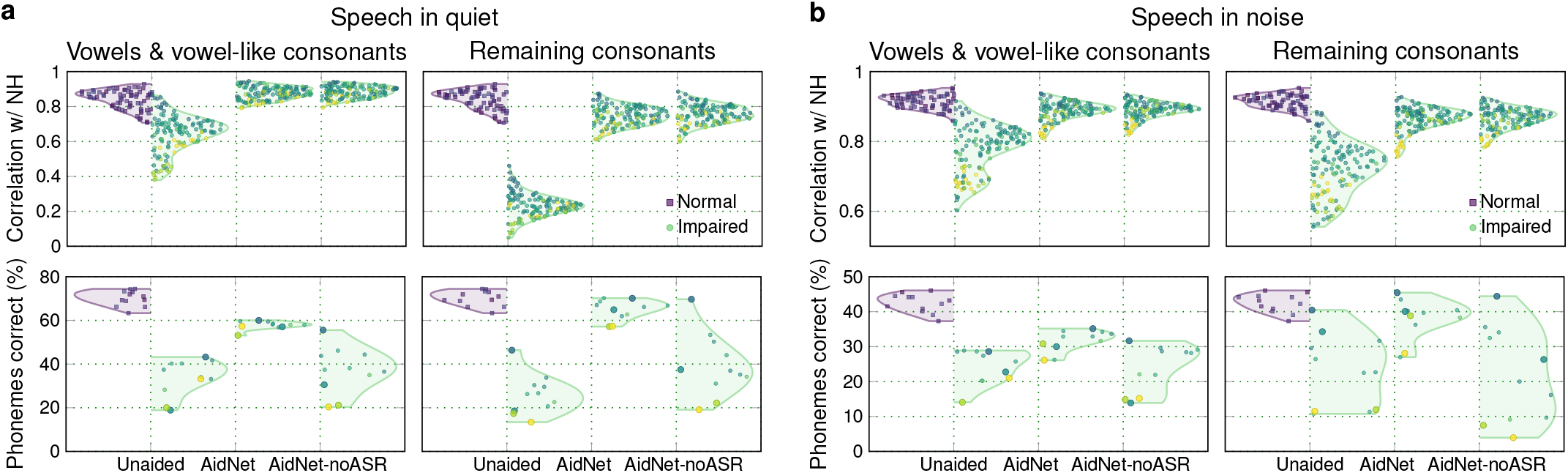
Restoration results without the inclusion of phoneme recognition loss in AidNet training. Restoration results are shown for the AidNet models trained with and without the phoneme recognition loss of the ASR model. The expectation over counts of 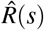 was taken for each ICNet model, and the metrics were computed after applying MCA alignment and flattening the responses. For phoneme recognition, larger markers indicate animals that were not included in the training of the ASR backend. We found that phoneme recognition performance was similar across animals with the same hearing status regardless of whether they were part of the training (green circles), both before and after processing sound. **a**. Results for the vowel and consonant parts of speech presented at 50 dB SPL in quiet. **b**. Results for speech mixed with restaurant noise at 0 dB SNR and presented at 65 dB SPL.

**Supplementary Fig. 5.**
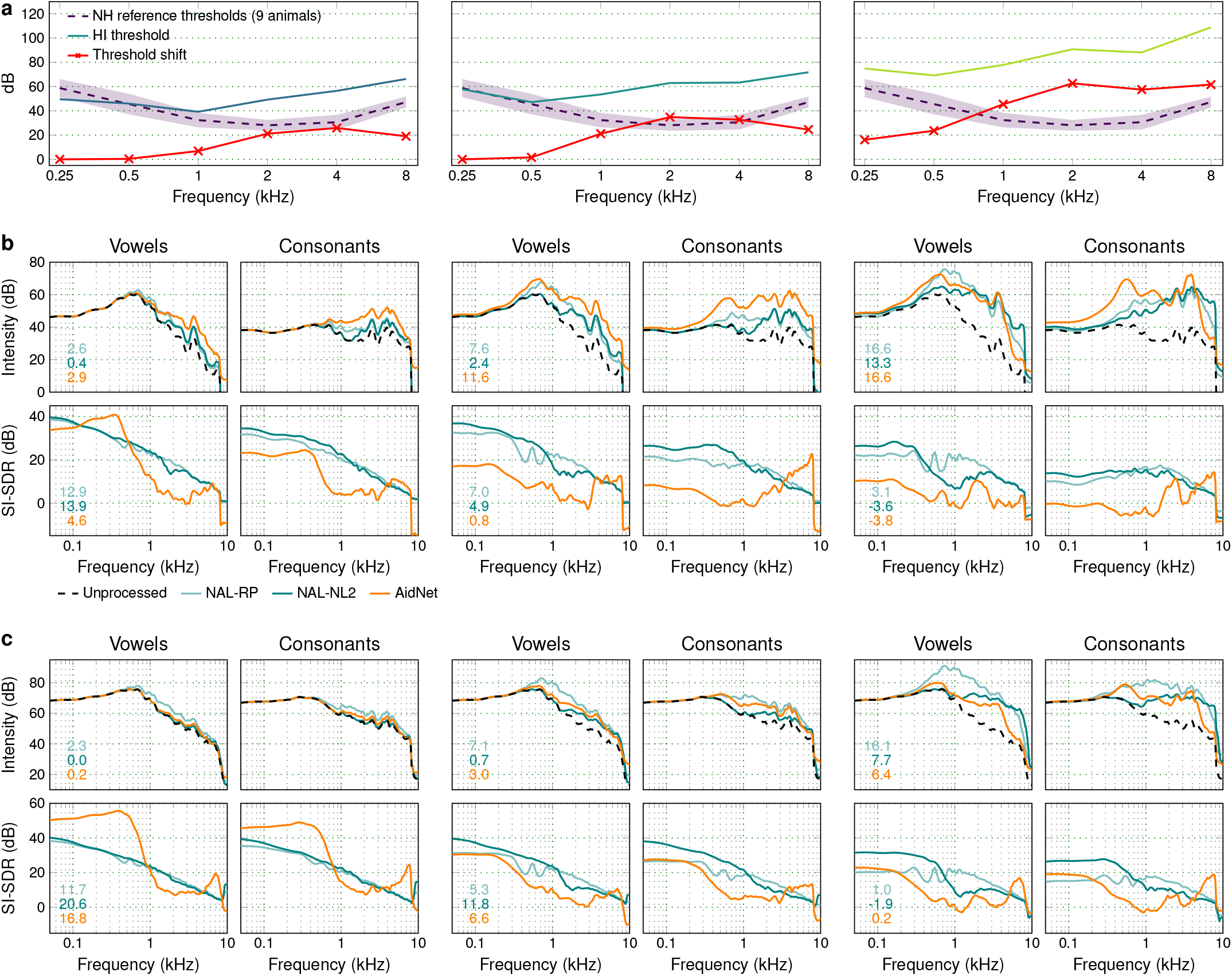
Example sound processing provided by AidNet. Sound processing results provided by 3 example AidNet models. Results of each column correspond to 3 HI animals with different hearing loss profiles. **a**. Hearing thresholds of the 3 animals that were used to fit the baseline hearing aid strategies. The threshold shift corresponds to the difference between each individual HI threshold and the NH average (clipped at zero). **b**. The sound processing that was provided for speech in quiet presented at 50 dB SPL is shown for the parts of speech that contained either vowels and vowel-like consonants (Vowels) or the remaining consonants (Consonants). The top rows show the frequency spectra before and after processing, while the bottom rows show the scale-invariant signal-to-distortion ratio (SI-SDR) after processing. The numbers correspond to the average gain and SI-SDR computed across the entire sound, with higher gain values indicating higher-amplitude sounds and lower SI-SDR values indicating a larger (scale-invariant) difference from the original sound. **c**. Sound processing results for speech in noise at 65 dB SPL and 0 dB SNR, presented as in **b**.

**Supplementary Fig. 6.**
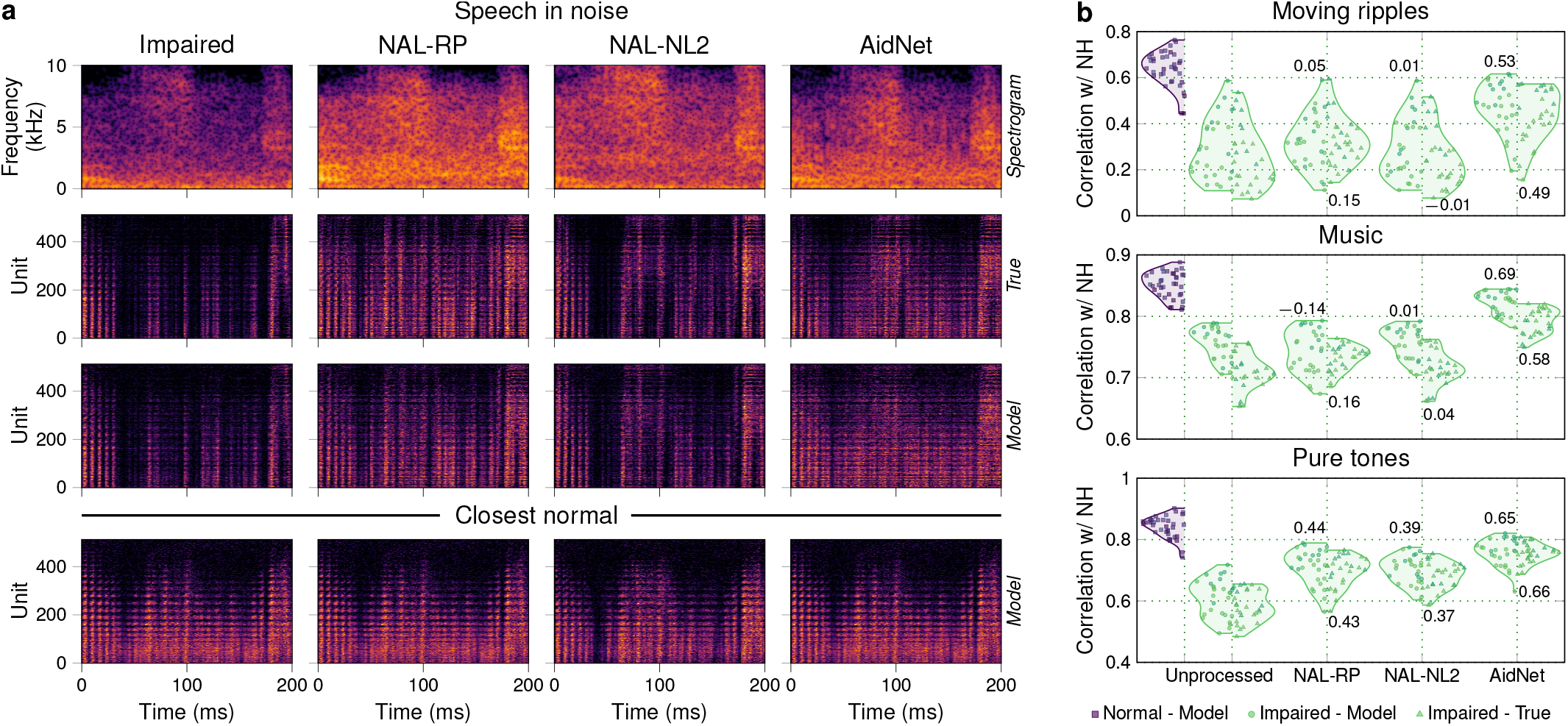
AidNets outperform the state of the art in vivo. **a**. Example spectrograms and responses to speech at 70 dB SPL and mixed with noise at 0 dB SNR, presented as in Fig. 4b. **b**. Restoration results for additional sounds that were not part of training, presented as in Fig. 4c. See also Supplementary Table 2.

**Supplementary Fig. 7.**
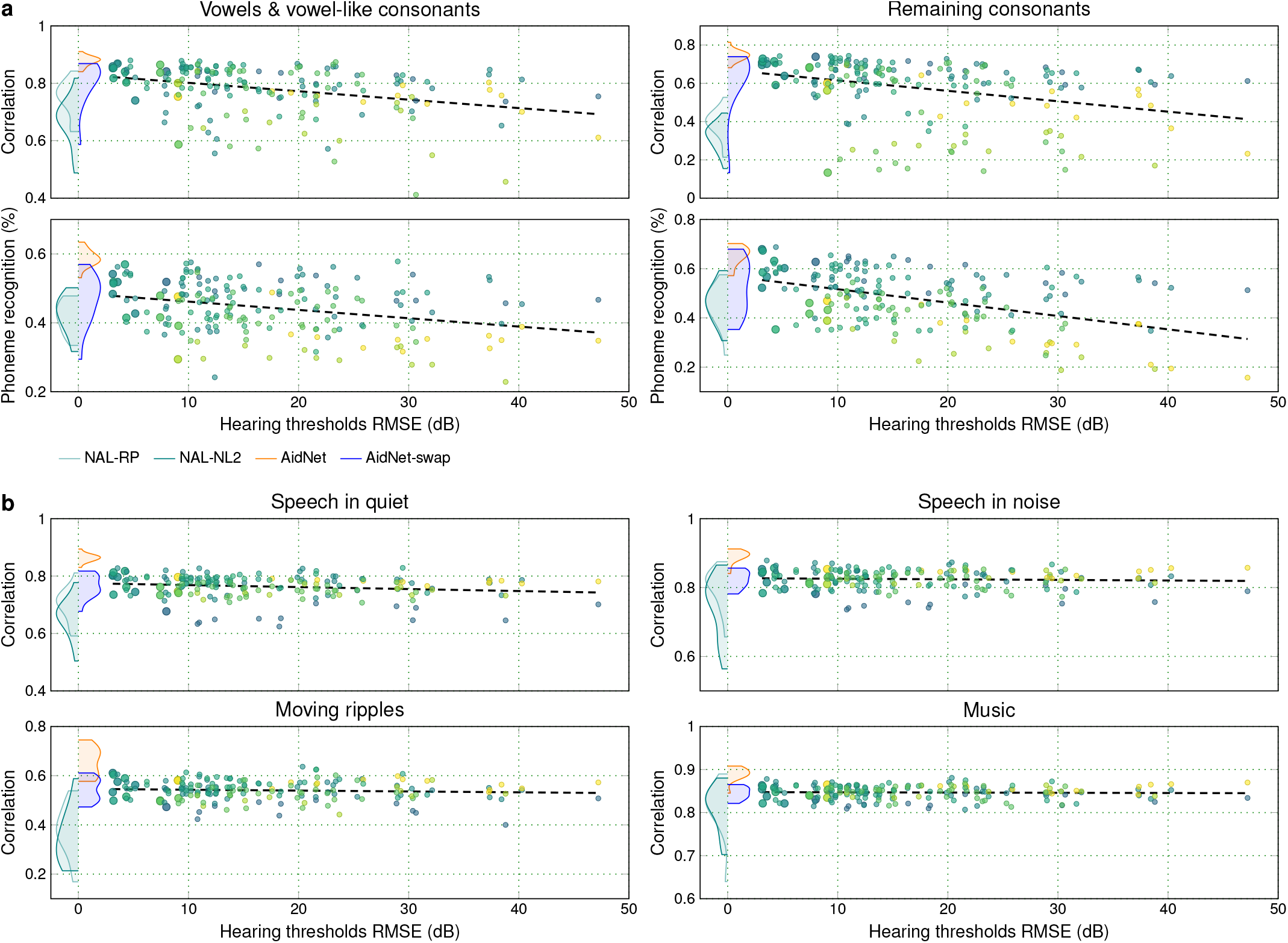
Full AidNet swap results for all sounds and animals. Simulated restoration results for 14 HI animals fitted with each of the AidNets of the other 13 animals are shown as a function of the difference in hearing thresholds between each pair of animals. **a**. Results for speech in quiet presented at 50 dB SPL (as in Fig. 3b, but averaged across NH animals for each HI animal and with the addition of the 3 HI animals used for the in vivo validation; see Supplementary Table 4). The results are shown separately for vowels and vowel-like consonants and the remaining consonants for both correlation and phoneme recognition. The dashed line corresponds to a linear fit to all data points. Larger markers indicate the results for each HI animal using the AidNet corresponding to the animal with the closest hearing threshold match (as in the AidNet-swap condition in Fig. 4d) and were used to compute the distributions in blue. The remaining distributions were computed in the same way from the fully individualized results (as in Fig. 3b, but averaged across NH animals for each HI animal). **b**. Correlation results for the four main evaluation sounds of Fig. 4d are presented as in **a**.

**Supplementary Fig. 8.**
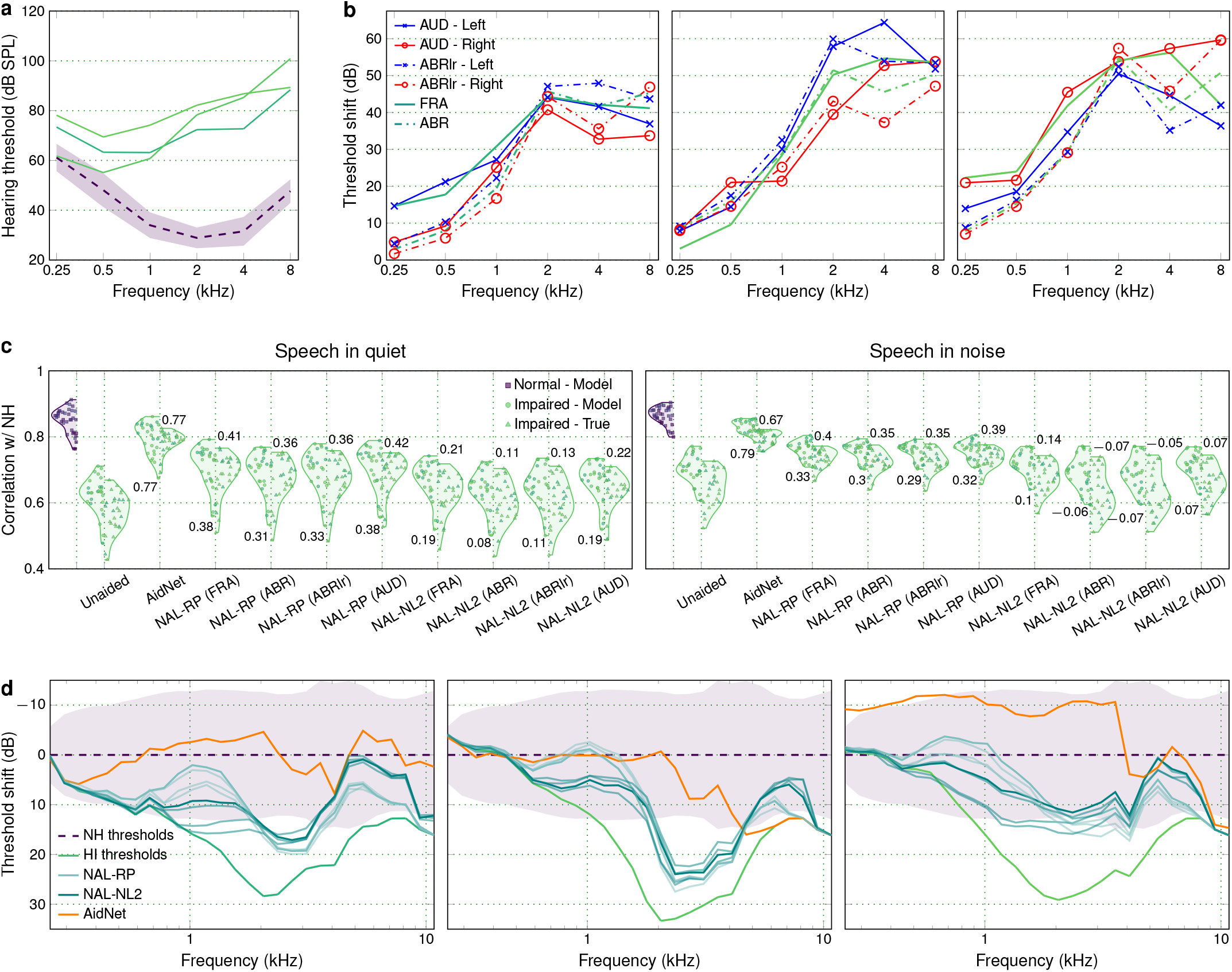
Comparison of different hearing threshold estimations and fittings. For each of the HI animals that were part of the in vivo validation, hearing thresholds were estimated from either IC activity or ABRs, and for either both ears or each ear separately. The effect of the different hearing threshold estimations was then tested by using the thresholds to obtain a different fitting for each of the benchmark hearing aid strategies before presenting the amplified sounds. **a**. Hearing thresholds of the 3 HI animals derived from the neural responses of both hemispheres to pure tones presented diotically (as in Fig. 1d). **b**. Threshold shifts were obtained for the 3 HI animals using four different hearing threshold estimates and were used to derive the gain prescriptions for each benchmark strategy. Solid lines correspond to thresholds estimated from IC neural activity, while dashed-dotted lines correspond to thresholds estimated from ABRs. Blue and red lines correspond to the thresholds of the left and right ear, respectively, obtained either from ABRs (ABRlr) or neural thresholds (AUD). Green lines correspond to thresholds estimated from both hemispheres (FRA; as in **a**) or the average between the left and right ABR thresholds (ABR), respectively. **c**. Effect of the different gain prescriptions on neural restoration for speech in quiet and in noise. Correlation was computed from real and predicted (sampled) responses averaged across 10 trials, after alignment with MCA and flattening across channels. The numbers indicate the average restoration index achieved by each processing strategy for recorded and simulated responses, with the FRA gain prescription (using the hearing thresholds of **a**) yielding the best results. **d**. Threshold shift as a function of frequency was estimated from the responses of all units to tones. Each line shows the average threshold shift of all units (relative to the mean of all units from 9 NH animals) derived from the frequency response areas at each tonal frequency after smoothing with a 2-D Gaussian window (standard deviation of 9 dB on the level dimension and 0.2 octaves on the frequency dimension). The shaded region indicates the standard deviation of the hearing thresholds of the 9 NH animals. For the two standard amplification strategies, the solid lines correspond to the FRA gain prescriptions (using the hearing thresholds of **a**) and the semi-transparent lines correspond to the remaining gain prescriptions corresponding to the remaining hearing threshold estimates of **b**.

**Supplementary Table 1.** In silico restoration results. Computed restoration indices for the results of Fig. 3b-c. The numbers correspond to the median and 95% confidence intervals (over bootstrap samples) of the restoration indices (*ρ*_*NH,HA*_ − *ρ*_*NH,HI*_)*/*(*ρ*_*NH*_ − *ρ*_*NH,HI*_) and (*PC*_*HA*_ − *PC*_*HI*_)*/*(*PC*_*NH*_ − *PC*_*HI*_) computed for the distributions of correlation and phoneme recognition, respectively, for 12 NH and 11 HI animals. Results are shown separately for vowels and vowel-like consonants (Vowels) and the remaining consonants (Consonants). The *p* values correspond to the Bonferroni-corrected results of a Wilcoxon rank-sum test between the indicated conditions after averaging across NH animals (*n* = 11 HI animals).

| Correlation | Speech in quiet |  | Speech in noise |  |
| --- | --- | --- | --- | --- |
|  | Vowels | Consonants | Vowels | Consonants |
| NAL-RP | 0.41 (0.38 0.44) | 0.27 (0.24 0.30) | 0.13 (0.08 0.17) | 0.19 (0.14 0.22) |
| NAL-NL2 | 0.15 (0.13 0.17) | 0.16 (0.13 0.18) | -0.05 (-0.09 -0.03) | 0.02 (-0.01 0.04) |
| AidNet | 0.90 (0.86 0.94) | 0.84 (0.81 0.87) | 0.77 (0.73 0.80) | 0.74 (0.71 0.77) |
| AidNet vs. NAL-RP | $p = 1.4e-04$ | $p = 1.4e-04$ | $p = 1.4e-04$ | $p = 1.4e-04$ |
| AidNet vs. NAL-NL2 | $p = 1.4e-04$ | $p = 1.4e-04$ | $p = 1.4e-04$ | $p = 1.4e-04$ |
| Phoneme recognition | Speech in quiet |  | Speech in noise |  |
|  | Vowels | Consonants | Vowels | Consonants |
| NAL-RP | 0.25 (0.19 0.30) | 0.44 (0.34 0.52) | 0.22 (0.13 0.30) | 0.37 (0.24 0.48) |
| NAL-NL2 | 0.29 (0.22 0.37) | 0.48 (0.38 0.56) | 0.24 (0.12 0.34) | 0.18 (0.01 0.28) |
| AidNet | 0.81 (0.79 0.83) | 0.89 (0.84 0.93) | 0.62 (0.52 0.71) | 0.94 (0.70 1.31) |
| AidNet vs. NAL-RP | $p = 1.4e-04$ | $p = 1.4e-04$ | $p = 3.2e-04$ | $p = 3.2e-04$ |
| AidNet vs. NAL-NL2 | $p = 1.4e-04$ | $p = 1.4e-04$ | $p = 8.9e-04$ | $p = 1.4e-04$ |

**Supplementary Table 2.** In vivo restoration results. Computed restoration indices for the results of Fig. 4c and Supplementary Fig. 6. The numbers correspond to the median and 95% confidence intervals (over bootstrap samples) of the restoration index (*ρ*_*NH,HA*_ − *ρ*_*NH,HI*_)*/*(*ρ*_*NH*_ − *ρ*_*NH,HI*_) computed for the correlation distributions of simulated (Model) and real (True) activity, for 9 NH and 3 HI animals. The small sample size (*n* = 3 HI animals) was not sufficient for formal statistical testing, but the AidNet restoration indices were always higher than the NAL-RP and NAL-NL2 restoration indices across all sounds and animals.

| Model | Speech in quiet | Speech in noise | Moving ripples | Music | Pure tones |
| --- | --- | --- | --- | --- | --- |
| NAL-RP | 0.38 (0.35 0.46) | 0.36 (0.30 0.47) | 0.05 (-0.02 0.16) | -0.14 (-0.23 0.02) | 0.44 (0.38 0.51) |
| NAL-NL2 | 0.19 (0.16 0.24) | 0.11 (0.08 0.15) | 0.01 (0.01 0.01) | 0.01 (-0.01 0.02) | 0.39 (0.33 0.45) |
| AidNet | 0.78 (0.68 0.89) | 0.80 (0.79 0.83) | 0.53 (0.48 0.62) | 0.69 (0.69 0.72) | 0.65 (0.63 0.72) |
| True | Speech in quiet | Speech in noise | Moving ripples | Music | Pure tones |
| NAL-RP | 0.40 (0.38 0.45) | 0.45 (0.41 0.49) | 0.15 (0.08 0.23) | 0.16 (0.11 0.23) | 0.43 (0.37 0.51) |
| NAL-NL2 | 0.20 (0.19 0.24) | 0.15 (0.13 0.17) | -0.01 (-0.02 0.00) | 0.04 (0.04 0.05) | 0.37 (0.34 0.43) |
| AidNet | 0.75 (0.73 0.78) | 0.68 (0.63 0.73) | 0.49 (0.44 0.55) | 0.58 (0.53 0.64) | 0.66 (0.61 0.72) |

**Supplementary Table 3.**
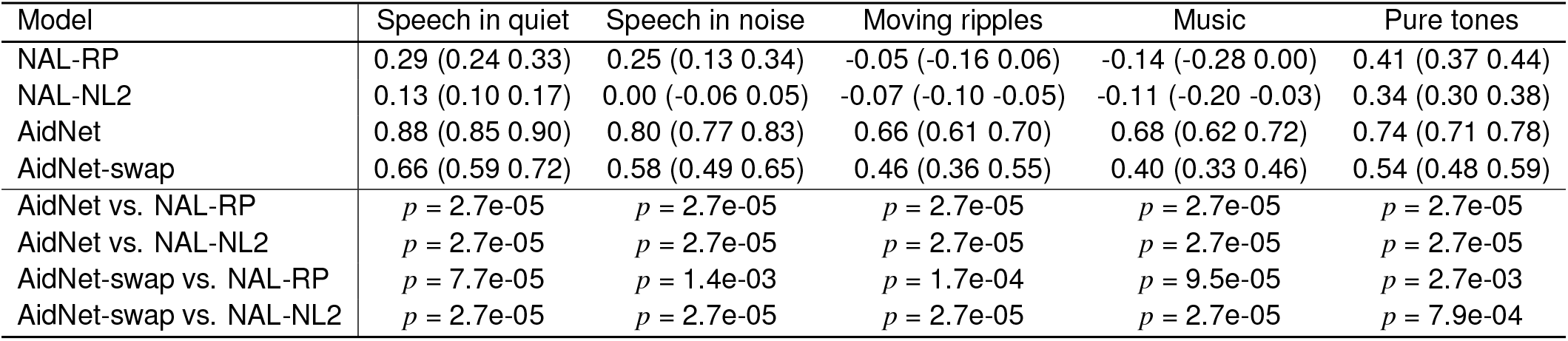
Simulated restoration results across all animals and sounds. Computed restoration indices for the results of Fig. 4d. The numbers correspond to the median and 95% confidence intervals (over bootstrap samples) of the restoration index (*ρ*_*NH,HA*_ − *ρ*_*NH,HI*_)*/*(*ρ*_*NH*_ − *ρ*_*NH,HI*_) computed for the correlation distributions of simulated activity, for 9 NH and 14 HI animals. The *p* values correspond to the Bonferroni-corrected results of a Wilcoxon rank-sum test between the indicated conditions after averaging across NH animals (*n* = 14 HI animals).

| Model | Speech in quiet | Speech in noise | Moving ripples | Music | Pure tones |
| --- | --- | --- | --- | --- | --- |
| NAL-RP | 0.29 (0.24 0.33) | 0.25 (0.13 0.34) | -0.05 (-0.16 0.06) | -0.14 (-0.28 0.00) | 0.41 (0.37 0.44) |
| NAL-NL2 | 0.13 (0.10 0.17) | 0.00 (-0.06 0.05) | -0.07 (-0.10 -0.05) | -0.11 (-0.20 -0.03) | 0.34 (0.30 0.38) |
| AidNet | 0.88 (0.85 0.90) | 0.80 (0.77 0.83) | 0.66 (0.61 0.70) | 0.68 (0.62 0.72) | 0.74 (0.71 0.78) |
| AidNet-swap | 0.66 (0.59 0.72) | 0.58 (0.49 0.65) | 0.46 (0.36 0.55) | 0.40 (0.33 0.46) | 0.54 (0.48 0.59) |
| AidNet vs. NAL-RP | $p = 2.7\text{e-}05$ | $p = 2.7\text{e-}05$ | $p = 2.7\text{e-}05$ | $p = 2.7\text{e-}05$ | $p = 2.7\text{e-}05$ |
| AidNet vs. NAL-NL2 | $p = 2.7\text{e-}05$ | $p = 2.7\text{e-}05$ | $p = 2.7\text{e-}05$ | $p = 2.7\text{e-}05$ | $p = 2.7\text{e-}05$ |
| AidNet-swap vs. NAL-RP | $p = 7.7\text{e-}05$ | $p = 1.4\text{e-}03$ | $p = 1.7\text{e-}04$ | $p = 9.5\text{e-}05$ | $p = 2.7\text{e-}03$ |
| AidNet-swap vs. NAL-NL2 | $p = 2.7\text{e-}05$ | $p = 2.7\text{e-}05$ | $p = 2.7\text{e-}05$ | $p = 2.7\text{e-}05$ | $p = 7.9\text{e-}04$ |

**Supplementary Table 4.** Animals used in the study. Recordings from 12 animals with normal hearing and 14 animals with hearing loss were included in total. The table shows the animals used for the analyses in each of the figures.

| Animal ID | Hearing | Dataset | Fig. 1c | Fig. 2b-c,<br>Fig. 3b-c,<br>SFig. 3-4 | Fig. 4c,<br>SFig. 6b,<br>SFig. 8 | Fig. 4d,<br>SFig. 7 | SFig. 5 |
| --- | --- | --- | --- | --- | --- | --- | --- |
| 240219 | Normal | Full | Yes | Yes | Yes | Yes | No |
| 240220 | Normal | Full | Yes | Yes | Yes | Yes | No |
| 240227 | Normal | Full | Yes | Yes | Yes | Yes | No |
| 240229 | Normal | Full | Yes | Yes | Yes | Yes | No |
| 240305 | Normal | Full | Yes | Yes | Yes | Yes | No |
| 240307 | Normal | Full | Yes | Yes | Yes | Yes | No |
| 240312 | Normal | Full | Yes | Yes | Yes | Yes | No |
| 240618 | Normal | Full | Yes | Yes | No | No | No |
| 240620 | Normal | Full | Yes | Yes | No | No | No |
| 240625 | Normal | Full | Yes | Yes | No | No | No |
| 231122 | Normal | Full | No | Yes | Yes | Yes | No |
| 231123 | Normal | Full | No | Yes | Yes | Yes | No |
| 250715 | Impaired | Full | Yes | Yes | No | Yes | No |
| 250410 | Impaired | Abridged | Yes | Yes | No | Yes | Yes |
| 240702 | Impaired | Full | Yes | Yes | No | Yes | No |
| 240514 | Impaired | Full | Yes | Yes | No | Yes | No |
| 240423 | Impaired | Full | Yes | Yes | No | Yes | Yes |
| 240709 | Impaired | Full | Yes | Yes | No | Yes | No |
| 231222 | Impaired | Full | Yes | Yes | No | Yes | No |
| 250424 | Impaired | Abridged | Yes | Yes | No | Yes | No |
| 250408 | Impaired | Full | Yes | Yes | No | Yes | Yes |
| 250401 | Impaired | Full | Yes | Yes | No | Yes | No |
| 231213 | Impaired | Full | No | Yes | No | Yes | No |
| 250924 | Impaired | Abridged | No | No | Yes | Yes | No |
| 250917 | Impaired | Abridged | No | No | Yes | Yes | No |
| 250909 | Impaired | Abridged | No | No | Yes | Yes | No |

## Notes

https://github.com/fotisdr/AidNet_example

https://doi.org/10.5281/zenodo.18407090

https://doi.org/10.5281/zenodo.19924033

### Summary of Updates

Main Figure updated, main text updated, new Supplementary Figures and Tables added

